# LIN28B is a post-transcriptional regulator of Wnt signalling in myeloid leukemia cells and dual targeting promotes cytotoxicity

**DOI:** 10.64898/2026.01.25.698240

**Authors:** O Sevim, M Wagstaff, RE Ling, A Goff, D Palmer, H Park, Katharine Hills, L Castellano, SF Newbury, A Roy, M Hartmann, BP Towler, RG Morgan

## Abstract

Wnt/β-catenin signalling is dysregulated across hematological malignancies but lacks effective targeting agents. Previously, we discovered that β-catenin interacts with RNA-binding proteins (RBP) and RNAs, indicating putative post-transcriptional roles in leukemia cells. The most frequent RBP-binding motif amongst β-catenin-bound mRNAs was the GGAG motif targeted by the oncofetal miRNA-regulating RBP; LIN28B. LIN28B is a critical regulator of fetal hematopoiesis often reactivated in rare myeloid neoplasms including juvenile myelomonocytic leukemia (JMML). In this study we investigated the molecular crosstalk between LIN28B and β-catenin in hematopoietic cells for the first time and examined dual targeting capacity. β-Catenin:LIN28B interaction was detected in both myeloid and lymphoid cell lines, and in primary human CD34^⁺^ fetal-liver derived HSCs. LIN28B regulated Wnt/β-catenin signalling through post-transcriptional modulation of *LEF1* via a *let7* miRNA-mediated mechanism. Single cell RNA sequencing data revealed *LIN28B* and *LEF1* correlation within abnormal JMML HSPCs enriched in high-risk hypermethylated subgroups characterised by oncofetal gene expression. miRNA sequencing of β-catenin- and LIN28B-depleted myeloid cells revealed shared miRNA targets, and dual pharmacological targeting preferentially reduced myeloid cell line viability. This study reveals Wnt/LIN28B molecular crosstalk in leukaemia cells which could represent a novel therapeutic axis in rare myeloid neoplasms harbouring β-catenin dysregulation and reactivation of LIN28B.

## Introduction

Canonical Wnt signalling is a transduction cascade critical for normal embryonic, fetal and adult development, and heavily dysregulated in human cancers. The pathway regulates several key adult tissue stem cell systems including the gut, breast, hair follicle, liver and blood.^1^ In particular, Wnt/β-catenin signalling plays a vital role in regulating the self-renewal^2^ and differentiation^3^ capacity of hematopoietic stem/progenitor cells (HSPC) in a dose-dependent fashion^4^ in both fetal and adult phases of hematopoiesis.^5^ Aberrant Wnt/β-catenin signalling has been reported across a plethora of hematological malignancies, where its activity contributes to the emergence, maintenance and drug resistance of leukemia stem cells (LSC).^6–10^ Despite extensive knowledge of deregulated Wnt signalling in human cancer, effective β-catenin targeting strategies remain absent from the clinic.

Previous approaches to targeting Wnt/β-catenin signalling have shown promise with a number of agents reaching Phase I or II clinical trials predominantly in solid cancers,^11^ as exemplified by WNT974 (PORCN inhibitor preventing Wnt ligand palmitoylation and secretion).^12^ However, very few of these have been deployed in a hematological setting which might partly be explained by β-catenin’s divergent molecular interactions in this context. For example, the well-characterised *APC*, *AXIN* and *CTNNB1* mutations which lead to Wnt signalling hyperactivation and initiate solid tumours such as colorectal cancer, are seldom reported in AML. Furthermore, β-catenin’s well characterised adhesion function at the cell membrane in adherens junctions is likely not a prominent feature of fluid hematopoietic tissue. Despite extensive characterisation in epithelial tissues, β-catenin’s molecular interactions/functions within hematological cells are much less well-defined and recent reports in leukemia settings suggest novel partnerships such as that with the transcription factors IRF7 or Ikaros.^13,14^ Our previous exploration of the β-catenin interactome in myeloid cells revealed the substantial enrichment of RNA binding proteins (RBP)^15,16^ indicating a potential novel role for β-catenin in the post-transcriptional landscape of gene regulation in leukemia cells.^17^ Since this time we have characterised several β-catenin:RBP interactions including WT1,^18^ TOE1,^19^ and MSI2^20^, and revealed β-catenin associates with an mRNA network of Wnt signalling and metabolic transcripts in leukemia cells.^20^ This study also suggested that β-catenin’s interaction with RNA is likely indirect and probably mediated through its various RBP partners. One putative interacting partner identified from our previous β- catenin interactome study was the oncofetal expressed, and miRNA regulating, RBP; LIN28B.^15^

Like OCT4 and SOX2, LIN28B is a critical pluripotency factor maintaining primitive stem cells,^21^ through both its role as a suppressor of *let7* miRNA biogenesis,^22^ and a direct binder of partner mRNAs.^23^ In a normal hematopoietic setting, its expression is confined to fetal HSPCs,^24^ where it maintains stemness and suppresses differentiation through *let7-*Cbx2,^25^ *let7*-Hmga2,^26^ and IGF2BP3 axes.^27^ However, the reactivation of LIN28B is observed in rare subsets of myeloid neoplasms including JMML.^28,29^ Despite previous reports of β-catenin and LIN28/let7 crosstalk in several tissues including breast,^30^ intestine,^31^ and lung,^32^ there are no studies examining the relationship of these two proteins in a hematopoietic context. This study examined the relationship between β-catenin and LIN28B in hematological cells and evaluate any potential dual-targeting capacity in myeloid leukemia.

## Methods

More in depth methodology is provided in *Supplementary Information*.

### Cell culture and drug treatments

Myeloid cell line culture, and CHIR99021 treatment (Merk-Millipore, Gillingham, Dorset), were performed as previously described.^15^ LIN28-*let-7* interaction was inhibited using compound 1632 (C1632)^33^ (Merck-Millipore) or ln268 (MedChemExpress Europe, Sollentuna, Sweden). β-Catenin inhibition in myeloid cell lines was attempted with BC2059, MSAB, KYA1797K, LGK974, Pyrvinium Pamoate, or XAV-939 (all MedChemExpress Europe). Global transcription was inhibited for 0, 4 and 8 hours using 5 µg/ml Actinomycin D (Merck-Millipore).

### Primary samples

Cord blood and fetal samples were approved and collected from institutions/committees outlined in Supplementary Information, with CD34^+^ HSPC fractions isolated as previously described.^34^

### Lentivirus preparation and transduction

MISSION lentiviral shRNA plasmids and lentiviral overexpression plasmids were obtained from Merck-Millipore and VectorBuilder, respectively (see Supplemental Table S1 for details). Transfection and transduction procedures are outlined in supplementary methods.

### Immunoblotting

Performed as previously described,^15^ using primary antibodies listed in Supplemental Table S2 and method extended in Supplementary information.

### Co-immunoprecipitation (Co-IP)

Performed as previously described,^15^ with Protein G Dynabeads (Thermo Fisher Scientific) coupled to 8µg of LIN28B antibody (Cell Signalling Technology, 4196) or rabbit IgG (Cell Signalling Technology, 2729).

### RNA extraction, clean-up and RT-qPCR

Performed as previously described,^20^ using the forward and reverse primer sequences listed in Supplemental Table S3 and optimised to show efficiency between 90-110%.

### Motif Analysis

Our previous β-catenin RIP-seq analysis identified the 190 common and significantly enriched transcripts in K562 and HEL cells. The full-length mature sequence of the longest isoform of each of these were used in motif analysis using the STREME tool within the MEME Suite searching for motifs between 6-15nt and the model of control sequences as a background.^35^

### Primary JMML scRNA sequencing analysis

*LIN28B* and *LEF1* expression was interrogated across primary JMML patient scRNA-seq datasets from Hartmann et al,^36^ and Louka et al^37^ using approach outlined in supplementary methods.

### miRNA sequencing

Total RNA extraction was performed from 2×10^6^ K562 cells using the Quick-RNA™ Miniprep Kit (Cambridge Bioscience) as per manufacturer’s guidelines. Quality control of total RNA samples was done with the Agilent TapeStation 4150. RNA samples were miRNA sequenced by BGI Genomics (Shenzhen, China) including library preparation and sequencing. FASTQ files were pre-processed them in sRNAbench on the sRNAtoolbox platform,^38^ with the following parameters: miRbase release 22.1 (as miRNA annotation reference database), Homo sapiens (hsa) (as short name from miRbase), human (GRCh38_p13) (as species), “Provided reads are already trimmed” (as sequencing library protocol), the rest of parameters as defaults. Raw read counts obtained after preprocessing was used to determine upregulated or downregulated miRNAs with β-catenin or LIN28B knockdown versus control using DESeq2 (version 1.44.0) package in R software.^39^ Also, <u>tidyverse (version 2.0.0)</u> and ComplexHeatmap (version 2.20.0) packages were used for data manipulation and visualisation.^40^ All LIN28B and β-catenin depleted K562 miRNA sequencing data has been uploaded to ArrayExpress and can be accessed using identifier E-MTAB-16028.

### RNA immunoprecipitation (RIP) / Cross-linking immunoprecipitation (CLIP)

RIP and CLIP assays were done as previously described.^20^

### Flow cytometry

TCF/LEF activity was assessed using the BAR/fuBAR reporter system as previously.^20^ Cell viability was determined using 1µg/ml 7-AAD viability staining solution (Thermo Fisher Scientific) or the Zombie Violet Fixable Viability Kit (BioLegend, London, UK).

### Statistics

Experiments were performed with three experimental replicates unless otherwise stated and statistical tests performed using GraphPad Prism (version 10; GraphPad, Boston, MA). Statistical analyses were conducted using Student’s *t*-test or ANOVA, with a threshold of *p* < 0.05 used for statistical significance. Data are presented as mean ± 1 standard deviation unless otherwise stated.

## Results

### β-Catenin and LIN28B interact in myeloid leukemia cell lines and fetal HSPC

Our previous exploration of the β-catenin interactome in hematopoietic cells revealed a dense RBP network,^15^ and subsequent β-catenin RIP-seq of myeloid cells identified β-catenin enrichment with mRNAs encoding regulators of Wnt signalling pathway and metabolism.^20^ These studies also suggested β-catenin has little direct RNA binding capacity and so RNA association is likely mediated through its many canonical RBP partners.^15,16^ To shortlist the RBP partners most important for β-catenin’s mRNA interactions, we analysed the most frequently occurring RBP binding motifs amongst the 190 commonly and significantly enriched transcripts derived from β-catenin RIP-seq in K562 and HEL cells. From this analysis we identified the GGAG motif to be one of the most frequently enriched binding motifs (p=0.019) which is bound by the *let7* miRNA regulating and oncofetal expressed RBP; LIN28 (Figure 1A).^41^ LIN28B was identified as a putative β-catenin interacting protein in myeloid cells from our original interactome study,^15^ suggesting any β-catenin influence over post-transcription could be mediated through its LIN28B interaction.

**Figure 1.**
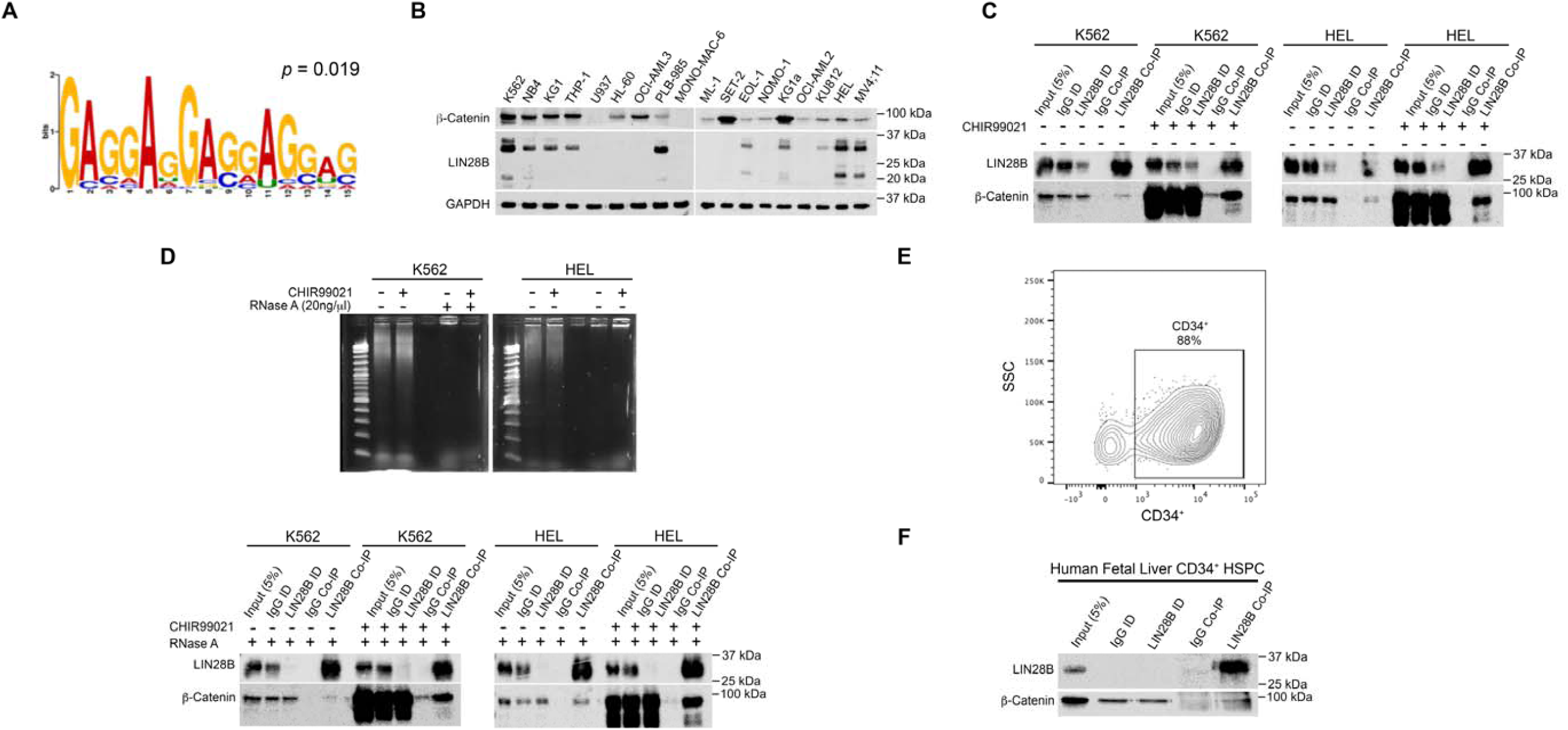
β-Catenin and LIN28B interact in myeloid cell lines and fetal HSPC. **(A)** RBP binding motif analysis from transcripts commonly and significantly enriched from β-catenin-RIP-seq performed in K562 and HEL cells.^20^ **(B)** Immunoblots showing β-catenin and LIN28B (long and short isoforms) expression across 18 myeloid leukemia cell lines. GAPDH was used as a loading control. Immunoblots show β-catenin level in LIN28B Co-IPs from K562 and HEL whole cell lysates following overnight ± 5μM CHIR99021 treatment. **(C)** Immunoblots showing β-catenin levels in LIN28B Co-IPs from K562 and HEL whole-cell lysates following overnight treatment ± 5μM CHIR99021. **(D)** Immunoblots showing β-catenin levels in LIN28B Co-IPs from K562 and HEL whole-cell lysates following ± overnight treatment 5μM CHIR99021 ± or 1hr pre-RNase A treatment (20µg/mL). **(E)** Flow-cytometric contour plot demonstrating the purity of human FL-CD34^⁺^ HSCs following FACS enrichment. **(F)** Immunoblot showing level of β-catenin and LIN28B from LIN28B Co-IPs executed from purified human FL-CD34^⁺^ HSCs. ID = immunodepleted lysate.

To interrogate this interaction further we examined the LIN28B and β-catenin expression across a panel of 18 myeloid cell lines previously screened for RBP/β-catenin expression.^19,20^ We observed variable expression of both the long and short LIN28B isoforms in approximately 50% of myeloid cell lines (Figure 1B), and co-expression of both LIN28B and β-catenin was detected in numerous cell lines which didn’t particularly align with any specific geno-or phenotypic features. Notably, the alternative LIN28 paralogue, LIN28A, was not detectable in any of the myeloid cell lines examined but was detected in the T47D breast cancer cell line, confirming antibody functionality (Supplementary Figure S1A). Based on this screen, we opted to validate the β-catenin:LIN28B interaction in the two cell lines used from the original TMT mass spectrometric analysis of the original β-catenin interactome study.^15^ As observed in Figure 1C, LIN28B was efficiently pulled-down from both K562 and HEL cells under both basal and Wnt stimulated conditions (CHIR99021 treated) and exhibited β-catenin interaction in both contexts. We also confirmed the β-catenin:LIN28B interaction in lymphoid SEM cells suggesting this interaction extends beyond a myeloid context (Supplementary Figure S1B). Given the association of both LIN28B^42^ and β-catenin^17^ with RNA, we wanted to rule out non-specific interaction of both proteins due to RNA co-occupancy. Complete digestion of RNA in input lysates prior to co-IP through RNase A treatment, had no impact on the β-catenin:LIN28B interaction (Figure 1D), suggesting the two molecules may be complexed together regardless of RNA presence. However, AlphaFold modelling suggests against a direct interaction between the two proteins suggesting other common intermediate proteins or structures could mediate the association (Supplementary Figure S1C).

Given LIN28B’s well characterised role as a regulator of fetal HSC emergence and development,^24–27^ a setting also requiring active Wnt/β-catenin signalling,^5^ we repeated the LIN28B Co-IP from human fetal liver (FL)-derived CD34^+^ HSPC. After confirming enrichment of FL-CD34^+^ HSPC to >85% purity (Figure 1E), and abundant levels of both proteins in input lysates, we confirmed β-catenin enrichment through LIN28B Co-IP (Figure 1F). Taken together, these findings identify LIN28B as a novel interacting partner of β-catenin in both myeloid and lymphoid cell lines, and primary human FL-CD34^+^ HSPCs.

### LIN28B modulates Wnt/β-catenin signalling output through LEF1 via a post-transcriptional mechanism

Given previous reports of LIN28B and Wnt signalling crosstalk in solid tissues,^30^ we assessed the degree and direction of crosstalk in leukemia cells. To examine this, we employed the β-catenin activated reporter (BAR) system adopted previously,^20^ to assess TCF/LEF activity upon LIN28B modulation. Upon confirmation of efficient LIN28B knockdown using shRNA (Figure 2A), we observed a ∼2-fold decrease in TCF/LEF activity in K562 cells upon Wnt signalling activation (Figure 2B and C). Reciprocally, ectopic LIN28B promoted a ∼2-fold increase in TCF/LEF activity under active Wnt signalling conditions (Figures D and E). Collectively, these results indicate that LIN28B is a positive regulator of Wnt/β-catenin signalling in myeloid cells.

**Figure 2.**
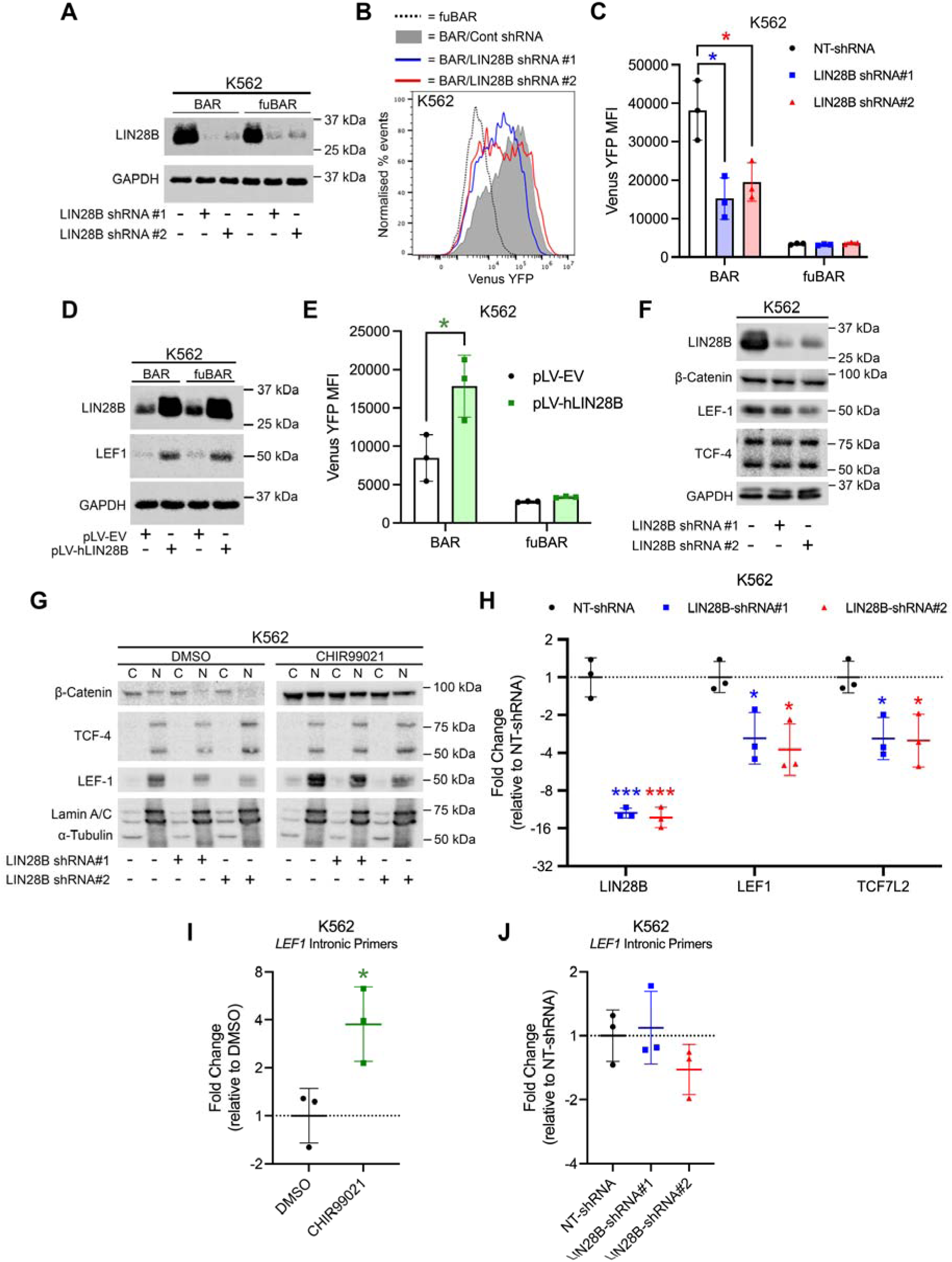
LIN28B regulates Wnt signalling through LEF-1 via a post-transcriptional mechanism. **(A)** Immunoblots showing LIN28B expression in K562 cells harbouring the BAR (β-catenin–activated reporter) or fuBAR (found unresponsive BAR; negative control) system with ± LIN28B shRNA. **(B)** Representative flow-cytometric histograms showing Venus Yellow Fluorescent Protein (YFP) intensity, reflecting TCF-dependent activity, in K562 cells with ± LIN28B shRNA following overnight treatment ± 5μM CHIR99021. The dashed histogram represents the fuBAR control, while the grey-filled histogram corresponds to non-targeting control shRNA. Blue and red histograms represent two LIN28B shRNAs. **(C)** Summary bar graph showing YFP MFI from the BAR/fuBAR system in K562 cells with ± LIN28B shRNA following overnight treatment ± 5μM CHIR99021. **D)** Immunoblot showing LIN28B and LEF-1 protein expression in K562 cells transduced with full-length human LIN28B cDNA (pLV-hLIN28B) versus an empty-vector (EV) control with GAPDH used as a loading control. **(E)** Summary bar graph showing YFP MFI from the BAR/fuBAR system in K562 cells ± pLV-hLIN28B following overnight treatment ± 5μM CHIR99021. **(F)** Immunoblots showing expression of LIN28B, β-catenin, LEF-1, and TCF4 in K562 cells with ± LIN28B shRNA. GAPDH was used as a loading control. **(G)** Immunoblots showing β-catenin, LEF-1, and TCF4 protein expression in cytoplasmic (C) and nuclear (N) fractions of K562 cells with ± LIN28B shRNA following ± 5μM CHIR99021. Lamin A/C and α-tubulin were used as fraction purity and loading controls. **(H)** Scatter plot showing fold changes in *LIN28B*, *LEF1*, *TCF7*, and *TCF7L2* mRNA levels in K562 cells with ± LIN28B shRNA, normalized to *GAPDH*. Summary graphs showing the fold change of immature *LEF1* mRNA levels assessed using primers spanning intronic regions of the *LEF1* gene in **(I)** K562 cells after overnight treatment with ± 5μM CHIR99021, and in **(J)** K562 cells with ± LIN28B shRNA. Data are presented as mean ± SD from three independent biological replicates. Statistical significance was assessed using Student’s t-test (\**p* < 0.05, \*\*\**p* < 0.001).

To investigate the underlying mechanism of LIN28B regulation of Wnt signalling output we first examined β-catenin expression and localization following LIN28B ablation. We observed no significant changes to overall β-catenin level (Figure 2F) or subcellular localization (Figure 2G). We examined the level of the downstream Wnt transcription factors LEF1 and TCF4 (*TCF7L2*) and observed a consistent reduction in both *LEF1* mRNA and protein levels in K562 cells following LIN28B depletion (Figure 2F, G, and H). Reciprocally, ectopic expression of LIN28B in K562 cells led to substantially increased LEF-1 protein levels (Figure 2D). These data indicate LIN28B positively regulates Wnt/β-catenin signalling activity through modulation of the downstream transcription factor LEF-1.

We next determined whether LIN28B regulation of LEF-1 occurred through transcriptional mechanisms since LIN28B has been linked with transcriptional regulation previously.^43^ To test this, we designed *LEF1* primers spanning intronic regions of *pre*-*LEF1* and assessed abundance following LIN28B depletion. As a control, we first validated these intronic primers in K562 cells treated with CHIR99021 to induce Wnt signalling activation since *LEF1* is a known Wnt signalling target gene,^44^ which confirmed a 4-fold increase compared to DMSO-treated cells (Figure 2I). Using these validated primers, we measured *LEF1* levels in LIN28B- depleted K562 cells and observed no change versus control NT-shRNA K562 cells (Figure 2J), suggesting regulation through post-transcriptional means.

### LIN28B and LEF1 are co-expressed in aberrant primary JMML subpopulations associated with high-risk disease

LIN28B expression is mainly confined to fetal tissue but is activated in several human malignancies. Indeed, JMML is reported to have one of the highest rates of LIN28B reactivation (∼50% cases) where it defines a novel fetal-like disease subgroup associated with risk.^28^ Given the scarcity of primary JMML patient samples available for analysis, we opted to interrogate existing bulk and single cell RNAseq patient datasets for potential *LIN28B* and *LEF1* relationship. Based on data from Louka et al., we subdivided JMML patients by low, medium and high *LIN28B* expression which revealed most similar co-expression with *LEF1* of the Wnt target genes analysed (Figure 3A).^37^ Further interrogation of the data revealed that *LIN28B*/*LEF1* co-expression was highest in the immunophenotypically aberrant ‘double positive’ (DP) CD45RA^+^CD90^+^ subpopulation that was identified within JMML samples in this study (Figure 3B and C). Building on the international consensus definition of risk-associated JMML DNA methylation epitypes (low methylation; LM, intermediate methylation; IM, and high methylation; HM),^45^ Hartmann and colleagues examined the JMML stem cell compartment and identified aberrant DP subpopulations predominantly in high-risk HM cases, whereas they were not detected in low-risk LM samples.^36^ Re-analysis of scRNA-seq data confirmed *LIN28B* expression in fetal but not postnatal HSPCs, largely coinciding with *LEF1* expression patterns (Figure 3D and E). *LIN28B*/*LEF*1 double-positive HSPCs were detected exclusively in intermediate- and high-risk JMML epitypes (IM and HM), whereas no double-positive cells were identified in low-risk LM JMML. Observed-to-expected analysis demonstrated enrichment of *LIN28B*+/*LEF1*+ cells in HM JMML, supporting a non-random association between *LIN28B* and *LEF1* detection (Figure 3E). Among *LIN28B*/*LEF1* double-positive HSPCs, *LIN28B* and *LEF1* expression levels were positively correlated, with the strongest association observed in HM JMML (Figure 3D). Finally, *LIN28B*- and *LEF1*-associated transcriptional programs derived from normal reference HSPCs showed strong correlations and elevated module scores in fetal and HM JMML HSPCs (Figure 3F and Supplementary Figure S2), consistent with coordinated activation of a shared oncofetal transcriptional state. In summary, these data indicate that a *LIN28B*:*LEF1* relationship could extend beyond immortalised cell lines and exists within aberrant subpopulations of adverse risk in primary JMML samples with an oncofetal signature.

**Figure 3.**
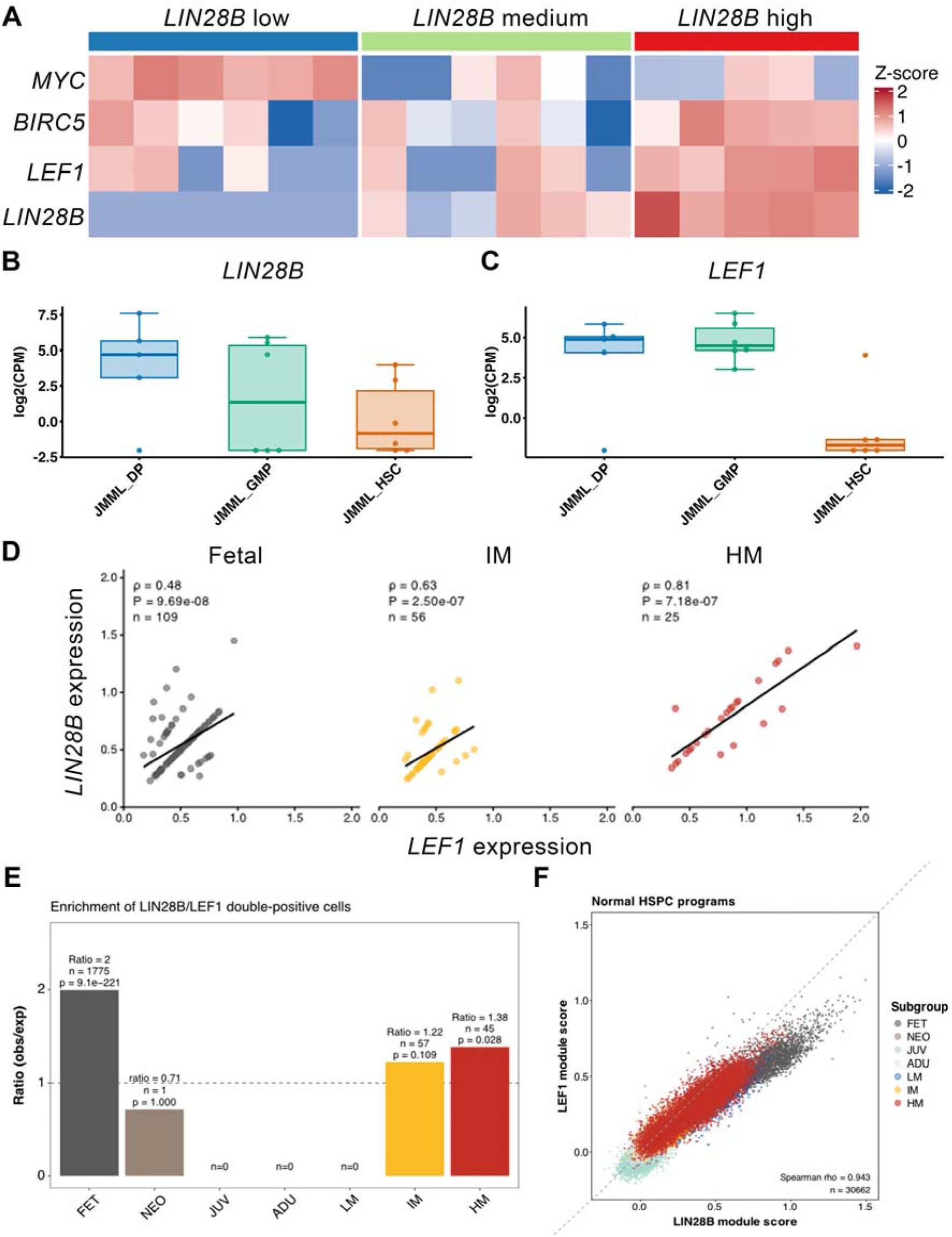
*LIN28B* and *LEF1* expression is correlated in aberrant JMML subpopulations enriched in high-risk disease. **(A)** Heat map showing the relative intensity (Z-score) of Wnt target gene expression (*MYC*, *BIRC5*, *LEF1*) across 17 primary JMML patient samples subdivided according to low (n=6), medium (n=6) or high (n=5) *LIN28B* expression. Box and whisker plots showing log2-transformed counts per million expression of **(B)** *LIN28B* and **(C)** *LEF1* across subpopulation of JMML aberrant double population (DP), JMML granulocyte-monocyte progenitors (GMP) and JMML HSC (n=6 per subgroup). **(D)** Scatter plots of *LIN28B* and *LEF1* expression in *LIN28B*+/*LEF1*+ fetal, IM JMML and HM JMML HSPCs. Only HSPCs with detectable expression of both transcripts were included. Each point represents one cell. Black lines indicate linear regression fits. Spearman correlation coefficients (ρ), two-sided P values and numbers of analysed cells (n) are shown for each subgroup. **(E)** Observed-to-expected ratios of *LIN28B*+/*LEF1*+ cells across developmental and JMML subgroups. The dashed line indicates the expected frequency under independent transcript detection (ratio = 1). P values were calculated using two-sided Fisher’s exact tests. Numbers above bars indicate the number of observed double-positive cells (n). **(F)** Scatter plots of normal HSPC-derived *LIN28B* and *LEF1* transcriptional programs. *LIN28B* and *LEF1* module scores were generated from gene signatures derived in normal reference HSPCs and calculated for individual normal and JMML HSPCs. Each point represents one cell and is coloured according to subgroup. Spearman correlation coefficient (ρ) and the number of analysed cells (n) are indicated.

### LIN28B regulates LEF1 via a post-transcriptional let7 miRNA-mediated mechanism

We hypothesised LIN28B could regulate LEF1 expression through direct transcript interaction since LIN28B is a well-known mRNA binding protein and can influence target expression/stability as evidenced with TP53^46^ and several splicing factors (*FUS/TLS*, *hnRNP F*, *TDP-43* and *TIA-1*).^41^ Adopting both LIN28B RIP and CLIP approaches from K562 cells, we first confirmed efficient enrichment of *LIN28B* mRNA as a positive control, since LIN28B is known to bind and autoregulate its own transcript (Supplementary Figure S3A-C).^41^ We also confirmed the significant enrichment of *LEF1* mRNA (Supplementary Figure S3A-C), however inhibiting global transcription using actinomycin D in LIN28B-depleted cells showed no marked difference from control cells (Supplementary Figure S3D), indicating LIN28B is unlikely to govern *LEF1* mRNA stability. We considered whether regulation of *LEF1* could occur through *let-7*-dependent mechanisms given the well-characterised role of LIN28B in the repression of *let-7* miRNA maturation.^22^ To first evaluate if LIN28B impacted *let-7* expression in myeloid cells we performed miRNA-seq in K562 cells with LIN28B shRNA (Figure 4A). We also examined miRNA expression in β-catenin depleted cells (Figure 4A) given its previous link regulating *let-7* expression in breast cancer cells,^30^ and to reveal any potential overlapping function with LIN28B. Clustering analysis revealed clear and distinct miRNA expression profiles across control, LIN28B and β-catenin shRNA samples (Figure 3B). Comparison of miRNA expression levels between both LIN28B-depleted and β-catenin–depleted cells relative to control (NT shRNA), revealed significantly upregulated and downregulated miRNAs in each condition (Figures 4C-H; see also Supplemental miRNA-seq data sheet). As expected, numerous *let-7* miRNAs, including *let-7d*, *let-7i* and *let-7f-1*, were significantly upregulated in response to LIN28B ablation (Figures 4C). In contrast, we observed no significantly differential *let-7* miRNA expression with β-catenin depletion in K562 cells (Figure 4D), suggesting β-catenin is not a major regulator of mature *let-7* miRNA expression in a myeloid setting. However, several miRNA species were differentially regulated in response to β-catenin modulation (Figure 4 and Supplemental Table S4), which were also affected upon LIN28B depletion (Figure 4 and Supplemental Table S5), suggesting novel miRNA regulating roles in hematopoietic cells which could be mediated through cooperation with LIN28B.

**Figure 4.**
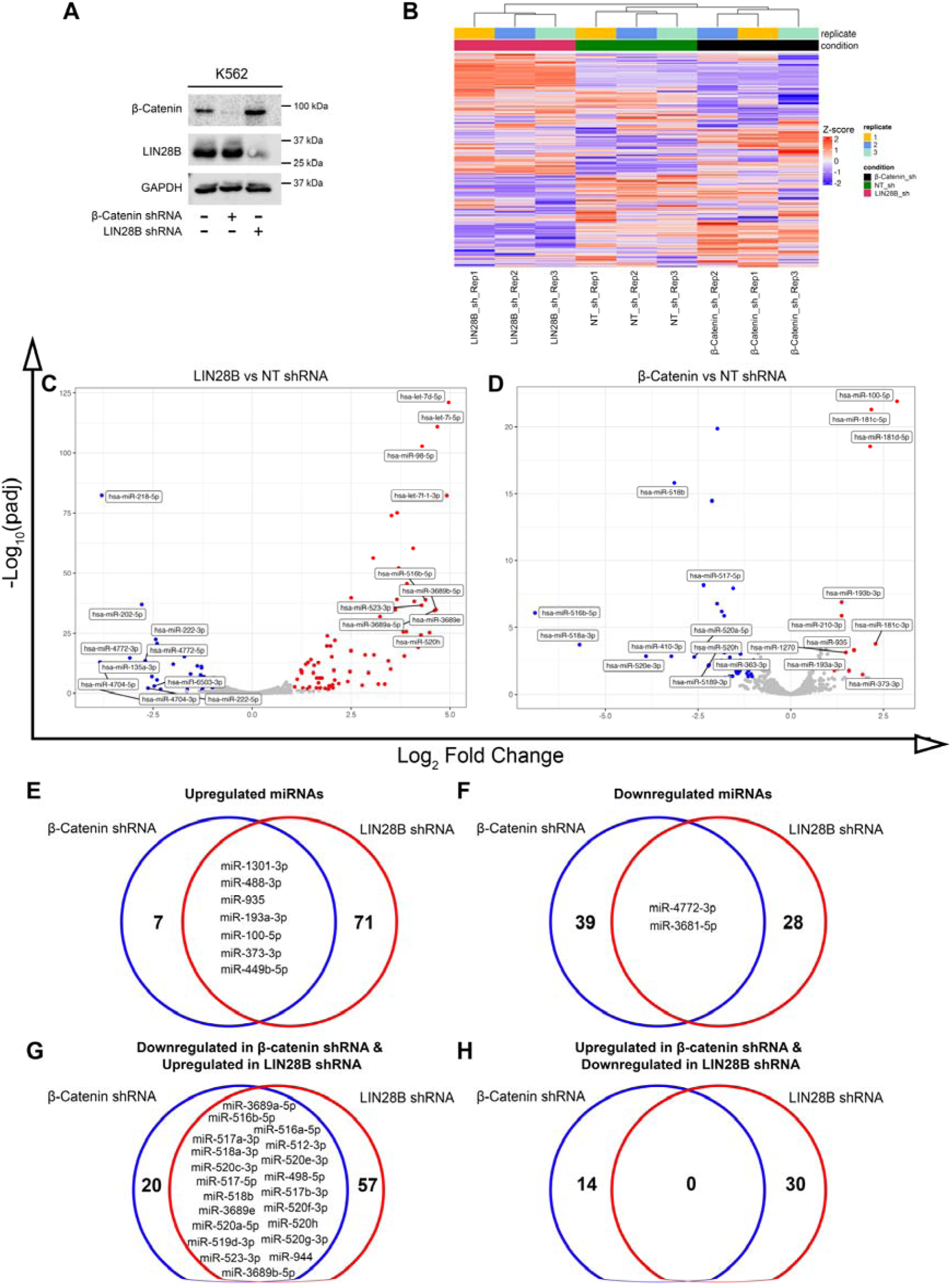
LIN28B and β-catenin regulate shared miRNA targets in myeloid cells. **(A)** Immunoblots show β-catenin and LIN28B expression levels in K562 cells transduced with NT, β-catenin, or LIN28B shRNA. GAPDH was used as a loading control. **(B)** A heatmap derived from scaled and normalized miRNA expression values in K562 cells with NT, β-catenin, or LIN28B shRNA is shown. The top dendrograms illustrate clustering of samples by replicate and condition. Volcano plots display significantly upregulated or downregulated miRNAs in K562 cells comparing **(C)** LIN28B shRNA vs. NT shRNA and **(D)** β-catenin shRNA vs. NT shRNA. Multiple testing correction was performed using the Benjamini–Hochberg method, and adjusted *p*-values are reported (*p*-adj). Red dots indicate significantly upregulated miRNAs (log_2_ fold change > 1 and *p*-adj ≤ 0.05), whereas blue dots denote significantly downregulated miRNAs (log_2_ fold change < –1 and *p*-adj ≤ 0.05). The highlighted points represent some of the top miRNAs in each comparison ranked by log2 fold change. Venn diagrams highlighting miRNAs commonly **(E)** upregulated or **(F)** downregulated in response to β-catenin or LIN28B shRNA in K562 cells. Venn diagrams highlighting potential antagonistically regulated miRNAs that are **(G)** downregulated by β-catenin shRNA but upregulated with LIN28B shRNA, and **(H)** upregulated by β-catenin shRNA but downregulated with LIN28B shRNA.

Given that multiple *let-7* miRNAs were induced following LIN28B suppression we opted to globally inhibit their expression, rather than interrogating individual *let-7* family members, as a means of evaluating *LEF1* regulation. For this we deployed C1632, a small molecule inhibitor (SMI) which blocks LIN28B:*let-7* interaction resulting in *let-7* induction.^33^ Following overnight treatment with two doses of C1632 we observed substantially reduced levels of LEF-1 protein (Figure 5A) and *LEF1* exonic mRNA (Figure 5B), in both K562 and HEL cells which was significantly depleted at 100µM, however intronic *LEF-1* remained unchanged (Figure 5C) confirming post-transcriptional, rather than transcriptional regulation, by C1632. We further observed LEF-1 reduction in response to a more recent and effective inhibitor of LIN28B:*let-7* interaction; ln268 (Supplemental Figure S4). Examination of TCF/LEF activity in C1632-treated cells, also revealed dose dependent depletion of Wnt signalling output in response to C1632 treatment in both K562 and HEL cells (Figure 5D). In summary, these findings indicate that LIN28B may regulate LEF-1 expression through a post-transcriptional *let-7* miRNA-mediated mechanism.

**Figure 5.**
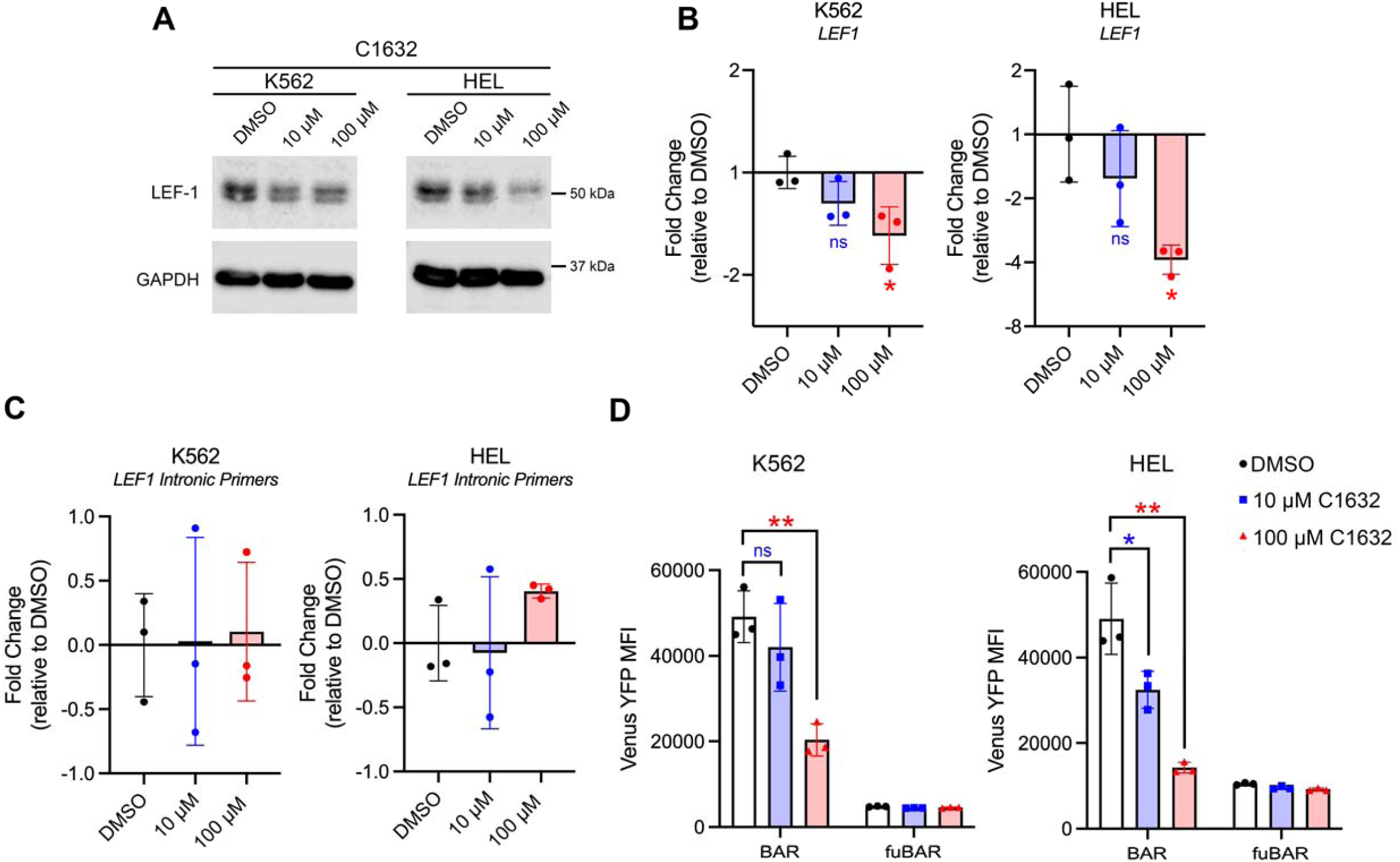
LIN28B regulates *LEF1* via a post-transcriptional *let7* miRNA-mediated mechanism. **(A)** Immunoblots showing the levels of LEF-1 protein in K562 and HEL cells following overnight treatment with 10μM or 100μM C1632. DMSO was used as a vehicle control. GAPDH served as a loading control. Bar graphs summarizing the fold change in *LEF1* mRNA levels in K562 and HEL cells after overnight treatment ± C1632 (10μM, 100μM) using primers to **(B)** exonic or (**C**) intronic regions, normalized to *GAPDH*. **(D)** Summary bar graphs showing MFI generated from BAR/fuBAR system in K562 and HEL cells following overnight treatment ± C1632 (10μM, 100μM) and ± 5μM CHIR99021. Data are presented as mean ± SD (standard deviation) from three independent biological replicates. Statistical significance was assessed using Student’s t-test (\**p* < 0.05; \*\**p* < 0.01; \*\**p* < 0.005; ns: *p* > 0.05).

### Dual β-catenin and LIN28B targeting decreases myeloid cell viability

We explored dual-targeting of the LIN28B:β-catenin axis given the molecular cooperation indicated from previous experiments. We deployed 100 µM C1632 and ln268 to target the LIN28B:let-7 interaction in K562 and HEL cells. We reasoned that compounds targeting β-catenin stability, rather than TCF/LEF or CBP/p300 interaction, were more applicable in this scenario given its interaction and subsequent miRNA regulation with LIN28B is likely post-transcriptional.

We first needed to identify an efficient β-catenin degrader in leukemia cells, since many SMIs were optimised in solid tissues where mechanisms of Wnt/β-catenin disruption differ from leukemia cells.^16,17^ To our surprise, most of the proposed β-catenin degraders were poorly effective at reducing β-catenin stability across a 24hrs exposure in myeloid cells including LGK974, MSAB, KYA1797K, XAV-939, and BC2059 (Tegatrabetan), and the β-catenin targeted PROTAC xStAx-VHLL (Supplementary Figure S5). These data suggest β-catenin stability is governed by additional mechanisms in leukemia cells. The only agent effective at reducing β-catenin stability at sub 50µM doses was pyrvinium pamoate (PP; Figure 6A and B) which is a Casein Kinase-1α (destruction complex component) agonist and an FDA-approved anthelmintic with anti-cancer activity, especially against MLL-rearranged AML.^47^ To evaluate a proof-of-concept approach to targeting LIN28:β-catenin, we deployed a dual-targeting strategy using both SMIs in K562 and HEL cells. As shown in Figure 6C and D, dual treatment of HEL cells with C1632 or ln268 and PP resulted in greater cell death than either drug alone. A similarly efficient dual-targeting effect was also observed in K562 cells (Figure 6E and F). Taken together, these data support the notion that dual β-catenin:LIN28B targeting could be an effective strategy in subtypes of myeloid leukemia like JMML where LIN28B is reactivated and β-catenin can be efficiently degraded.

**Figure 6.**
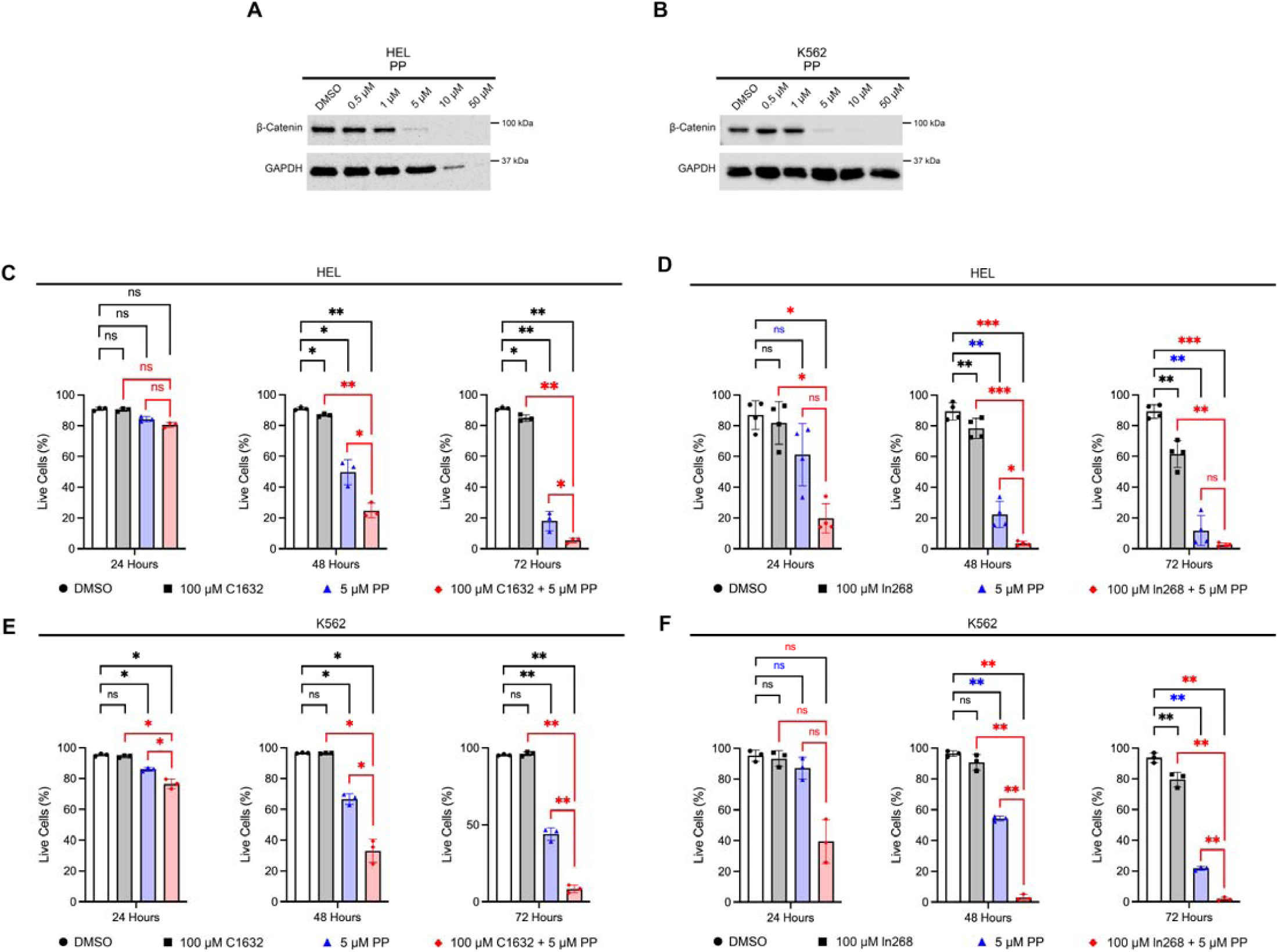
Dual β-catenin and LIN28B targeting decreases myeloid cell viability. Immunoblots showing the expression level of β-catenin protein in HEL **(A)** and K562 **(B)** cells following overnight treatment with different concentrations of Pyrvinium pamoate (PP). Bar graphs display the percentage of live cells as determined by 7AAD negative staining in HEL cells after treatment with **(C)** DMSO, 100μM C1632, 5μM PP and 100μM C1632 & 5μM PP, or **(D)** DMSO, 100μM ln268, 5μM PP and 100μM ln268 & 5μM PP for 24, 48 and 72 hours. Bar graphs display the percentage of live cells as determined by 7AAD negative staining in K562 cells after treatment with **(E)** DMSO, 100μM C1632, 5μM PP and 100μM C1632 & 5μM PP, or **(F)** DMSO, 100μM ln268, 5μM PP and 100μM ln268 & 5μM PP for 24, 48 and 72 hours. Data are presented as mean ± SD from three independent biological replicates. Statistical significance was assessed using Student’s t-test (\**p* < 0.05; \*\**p* < 0.01; \*\*\**p* < 0.001; ns: non-significant).

## Discussion

This is the first study to demonstrate a β-catenin and LIN28B interaction and molecular crosstalk. Our AlphaFold modelling suggests this interaction is indirect, with our data suggesting an RNA-independent association that may be part of a wider protein complex. Indeed, this is now the third RBP interaction with β-catenin that we’ve demonstrated has regulatory capacity over LEF-1 expression,^19,20^ and points to β-catenin constituting a wider post-transcriptional regulatory network that governs expression of this Wnt transcription factor. We’ve not been able to characterise the functional significance of the β-catenin:LIN28B interaction but our data suggests no capacity to regulate the stability/localisation of the other. Instead we speculate this interaction could impact miRNA expression since our miRNA-seq experiment identified co- and reciprocally regulated targets from LIN28B and β-catenin depletion, and cooperation between β-catenin and LIN28A in miRNA regulation has been reported in the expansion of breast cancer stem cells.^30^ This study also demonstrated that LIN28B reulgates LEF-1 expression, likely through a post-transcriptional mechanism. Crosstalk between Wnt/β-catenin signalling and LIN28 has been observed frequently in both normal development,^48^ and cancer,^31^ however this is the first report in a hematopoietic context and the first to implicate LEF-1 regulation. Our experiments using the LIN28B:*let7* inhibitor C1632 suggest this regulation is primarily post-transcriptional and potentially via miRNA regulation of *LEF1*. One previous study reports *let7* regulation of LEF-1 expression (via *let-7i-3p*),^49^ however there are numerous miRNAs reported to regulate *LEF1* expression including miR-34a,^50^ and miR-181a,^51^ which were amongst the most significantly increased miRNAs following LIN28B and β-catenin depletion, respectively, in our study.

LIN28B itself has an enigmatic and complex role in normal and malignant hematopoiesis. Expression is reportedly restricted to fetal hematopoiesis,^24^ where it regulates fetal HSC development through *let-7*–Hmga2,^26^ *let-7*-Cbx2,^25^ and IGF2BP3 axes.^27^ Active Wnt/β-catenin signalling also contributes to these early stages of hematopoiesis,^5^ and our finding that β-catenin interacts with LIN28B in purified human fetal liver CD34^+^ HSPCs raises the intriguing possibility of crosstalk/cooperation between these proteins in early hematopoietic development. Upon transition to adult hematopoiesis, LIN28B expression is rapidly downregulated (permitting mature *let-7* expression) with expression only reactivated in a small subset leukemias, or within the fetal-restricted programme necessary to expand adult HSCs ex vivo.^52^

There is evidence for LIN28B serving both tumour suppressive and oncogenic functions in malignant hematopoiesis. *In vivo* models of MLL-derived myeloid leukemogenesis (MLL-AF9 and MLL-ENL) have demonstrated that LIN28B expressing fetal progenitors have reduced transformation capacity, and only the reduction of LIN28B expression in these cell provides a leukemia-permissive environment.^53,54^ In myeloid neoplasms, LIN28B reactivation appears to have a pro-leukemic role contributing to differentiation arrest,^55^ growth/survival,^56^ and promotion of a stem-like transcriptional programme.^57^ In CML, LIN28B expression is enhanced in blast phase versus chronic phase, and promotes proliferation through a miR-181d-PDCD4 axis (miR-181d also upregulated in our b-catenin shRNA miRNA-seq) which involves a LIN28B DNA-binding function.^58^ Around half of JMML samples are reported to reactivate LIN28B which defines a fetal-like subgroup of JMML cases with adverse survival.^28,29^ The scarcity of JMML primary patient samples and the lack of *in vitro* JMML models prevented us investigating Wnt crosstalk further in this disease, however two recent JMML scRNAseq datasets indicated a significant LIN28B/LEF1 correlation,^36,37^ particularly evident in aberrant DP JMML subpopulations which are overrepresented in high-risk/HM subgroups. We speculate that LIN28B/LEF1 represents a developmental axis in normal fetal HSPC that is reactivated in high-risk/HM JMML subgroups conferring a fetal-like state.

Although this study has not elucidated the mechanistic role of the β-catenin:LIN28B interaction, we have shown that dual targeting of both molecules may have therapeutic merit in subsets of myeloid leukemia where they are both expressed. LIN28B inhibitors remain in their infancy and although C1632 has shown activity against several solid cancers,^59^ it’s not a particularly efficient compound requiring high doses such as those used in this study (50-100µM) to be effective. Several new LIN28 inhibitors have been developed, such as ln268,^60^ however we found high concentrations of this were also required. Regardless, if deployed properly they’ll exhibit a tolerable toxicity profile given LIN28B is not expressed in normal healthy adult tissues limiting on-target toxicity. We found both LIN28B inhibitors were minimally cytotoxic to normal human CB-derived CD34^+^ HSPC (Supplementary Figure S6).

Even if LIN28B inhibition can be refined, our study highlights that pharmacological β-catenin targeting in leukemia remains a formidable challenge. Many of the compounds proposed to reduce β-catenin level in solid tissues, including LGK974 (PORCN inhibitor), MSAB (β-catenin degrader), KYA1797K (Axin agonist), XAV-939 (Tankyrase inhibitor), BC2059 (Transducin β-Like 1 blocker), and xStAx-VHL (β-catenin PROTAC) were inefficient in leukemia cells, pointing to alternative modes of β-catenin stabilisation/regulation in myeloid leukemia yet to be defined. The only compound that showed any capacity for β-catenin degradation was PP which is an FDA-approved quinoline-derived cyanine dye and antihelminth drug that boosts Casein Kinase-1α activity, a critical destruction complex component providing the serine/threonine phosphorylation necessary for β-catenin’s subsequent ubiquitination and proteasomal degradation. In recent years, PP has emerged as an anti-tumour compound with activity against MLL-AML,^47^ however PP is not a specific Wnt/β-catenin inhibitor, also acting through mitochondrial disruption, PI3K/AKT inhibition, and UPR activation.^61^ Therefore, there remains an urgent and unmet need for specific and efficient β-catenin degraders in myeloid leukemia.

In summary, we’ve identified an interaction between β-catenin and LIN28B, and show that LIN28B can impact Wnt signalling activity in hematopoietic cells through the post-transcriptional regulation of LEF-1 expression. This study has demonstrated that dual targeting of β-catenin and LIN28B could represent a novel and effective strategy in rare myeloid neoplasms subsets where both proteins are reactivated.

## Author contributions

OS performed experiments, analysed data and co-wrote the manuscript. MW managed laboratory, performed experiments and provided experimental guidance. RL assisted with FL-CD34 HSC experiments whilst AG, DP, HP and KH generated contributing data. LC and SN provided reagents and experimental advice, while AR provided primary FL-CD34 HSCs and conceptual guidance. MH executed computational interrogation of primary JMML data. BT and RGM performed experiments, analysed data, co-wrote the manuscript and conceived/directed the study.

## Data availability

LIN28B and β-catenin depleted K562 miRNA sequencing data has been uploaded to ArrayExpress and can be accessed using identifier E-MTAB-16028. All other data featured in this manuscript are available from the corresponding author upon reasonable request

## Supporting information

Supplementary miRNA-seq dataset

## Acknowledgements

We thank all the technicians in the School of Life Sciences that maintain our laboratories and facilities. Thanks to the midwives and clinical research nurses at University Hospitals Sussex and East Sussex Healthcare NHS trusts for the collection of human umbilical cord blood. We are indebted to the patients and their families who gave consent for their samples to be used for our research.

## References

1 Maurice, M. M. & Angers, S. Mechanistic insights into Wnt-beta-catenin pathway activation and signal transduction. Nat Rev Mol Cell Biol 26, 371–388 (2025). 10.1038/s41580-024-00823-y

2 Reya, T. et al. A role for Wnt signalling in self-renewal of haematopoietic stem cells. Nature 423, 409–414 (2003).

3 Sheng, Y. et al. Activation of wnt/beta-catenin signaling blocks monocyte-macrophage differentiation through antagonizing PU.1-targeted gene transcription. Leukemia 30, 2106–2109 (2016). 10.1038/leu.2016.146

4 Luis, T. C. et al. Canonical wnt signaling regulates hematopoiesis in a dosage-dependent fashion. Cell Stem Cell 9, 345–356 (2011).

5 Ruiz-Herguido, C. et al. Hematopoietic stem cell development requires transient Wnt/beta-catenin activity. J Exp Med 209, 1457–1468 (2012). 10.1084/jem.20120225

6 Wang, Y. et al. The Wnt/beta-catenin pathway is required for the development of leukemia stem cells in AML. Science 327, 1650–1653 (2010).

7 Yeung, J. et al. beta-Catenin mediates the establishment and drug resistance of MLL leukemic stem cells. Cancer Cell 18, 606–618 (2010).

8 Dietrich, P. A. et al. GPR84 sustains aberrant beta-catenin signaling in leukemic stem cells for maintenance of MLL leukemogenesis. Blood 124, 3284–3294 (2014). 10.1182/blood-2013-10-532523

9 Fong, C. Y. et al. BET inhibitor resistance emerges from leukaemia stem cells. Nature 525, 538–542 (2015). 10.1038/nature14888

10 Perry, J. M. et al. Overcoming Wnt-beta-catenin dependent anticancer therapy resistance in leukaemia stem cells. Nat Cell Biol 22, 689–700 (2020). 10.1038/s41556-020-0507-y

11 Park, W. J. & Kim, M. J. A New Wave of Targeting ‘Undruggable’ Wnt Signaling for Cancer Therapy: Challenges and Opportunities. Cells 12 (2023). 10.3390/cells12081110

12 Rodon, J. et al. Phase 1 study of single-agent WNT974, a first-in-class Porcupine inhibitor, in patients with advanced solid tumours. Br J Cancer 125, 28–37 (2021). 10.1038/s41416-021-01389-8

13 Hosseini, A. et al. Perturbing LSD1 and WNT rewires transcription to synergistically induce AML differentiation. Nature 642, 508–518 (2025). 10.1038/s41586-025-08915-1

14 Cosgun, K. N. et al. Targeting beta-catenin degradation with GSK3beta inhibitors induces cell death in acute lymphoblastic leukemia. Nat Cancer 7, 150–168 (2026). 10.1038/s43018-025-01093-z

15 Morgan, R. G. et al. LEF-1 drives aberrant beta-catenin nuclear localization in myeloid leukemia cells. Haematologica 104, 1365–1377 (2019). 10.3324/haematol.2018.202846

16 Wagstaff, M., Coke, B., Hodgkiss, G. R. & Morgan, R. G. Targeting beta-catenin in acute myeloid leukaemia: past, present, and future perspectives. Biosci Rep (2022). 10.1042/BSR20211841

17 Sevim, O., Park, H. & Morgan, R. G. Post-transcriptional control of gene expression by beta-catenin: expanding the non-canonical ARMoury. Oncogene 44, 2453–2459 (2025). 10.1038/s41388-025-03470-5

18 Wagstaff, M. et al. Crosstalk between beta-catenin and WT1 signaling activity in acute myeloid leukemia. Haematologica 108, 283–289 (2023). 10.3324/haematol.2021.280294

19 Park, H. et al. TOE1 is a beta-catenin interacting protein regulating the proliferation of hematopoietic cells through PAK2 modulation. Stem Cell Reports 21, 102894 (2026). 10.1016/j.stemcr.2026.102894

20 Wagstaff, M. et al. beta-Catenin interacts with canonical RBPs including MSI2 to associate with a Wnt signalling mRNA network in myeloid leukaemia cells. Oncogene 44, 2490–2503 (2025). 10.1038/s41388-025-03415-y

21 Zhang, J. et al. LIN28 Regulates Stem Cell Metabolism and Conversion to Primed Pluripotency. Cell Stem Cell 19, 66–80 (2016). 10.1016/j.stem.2016.05.009

22 Piskounova, E. et al. Lin28A and Lin28B inhibit let-7 microRNA biogenesis by distinct mechanisms. Cell 147, 1066–1079 (2011). 10.1016/j.cell.2011.10.039

23 Tsanov, K. M. et al. LIN28 phosphorylation by MAPK/ERK couples signalling to the post-transcriptional control of pluripotency. Nat Cell Biol 19, 60–67 (2017). 10.1038/ncb3453

24 Yuan, J., Nguyen, C. K., Liu, X., Kanellopoulou, C. & Muljo, S. A. Lin28b reprograms adult bone marrow hematopoietic progenitors to mediate fetal-like lymphopoiesis. Science 335, 1195–1200 (2012). 10.1126/science.1216557

25 Wang, D. et al. Developmental maturation of the hematopoietic system controlled by a Lin28b-let-7-Cbx2 axis. Cell Rep 39, 110587 (2022). 10.1016/j.celrep.2022.110587

26 Copley, M. R. et al. The Lin28b-let-7-Hmga2 axis determines the higher self-renewal potential of fetal haematopoietic stem cells. Nat Cell Biol 15, 916–925 (2013). 10.1038/ncb2783

27 Wang, S. et al. Enhancement of LIN28B-induced hematopoietic reprogramming by IGF2BP3. Genes Dev 33, 1048–1068 (2019). 10.1101/gad.325100.119

28 Helsmoortel, H. H. et al. LIN28B overexpression defines a novel fetal-like subgroup of juvenile myelomonocytic leukemia. Blood 127, 1163–1172 (2016). 10.1182/blood-2015-09-667808

29 Helsmoortel, H. H. et al. LIN28B is over-expressed in specific subtypes of pediatric leukemia and regulates lncRNA H19. Haematologica 101, e240–244 (2016). 10.3324/haematol.2016.143818

30 Cai, W. Y. et al. The Wnt-beta-catenin pathway represses let-7 microRNA expression through transactivation of Lin28 to augment breast cancer stem cell expansion. J Cell Sci 126, 2877–2889 (2013). 10.1242/jcs.123810

31 Tu, H. C. et al. LIN28 cooperates with WNT signaling to drive invasive intestinal and colorectal adenocarcinoma in mice and humans. Genes Dev 29, 1074–1086 (2015). 10.1101/gad.256693.114

32 Zhang, T. et al. Glycogen synthase kinase-3beta promotes radiation-induced lung fibrosis by regulating beta-catenin/lin28 signaling network to determine type II alveolar stem cell transdifferentiation state. FASEB J 34, 12466–12480 (2020). 10.1096/fj.202001518

33 Roos, M. et al. A Small-Molecule Inhibitor of Lin28. ACS Chem Biol 11, 2773–2781 (2016). 10.1021/acschembio.6b00232

34 Morgan, R. G. et al. gamma-Catenin is expressed throughout normal human hematopoietic development and is required for normal PU.1-dependent monocyte differentiation. Leukemia 27, 2096–2100 (2013). 10.1038/leu.2013.96

35 Bailey, T. L. et al. MEME SUITE: tools for motif discovery and searching. Nucleic Acids Res 37, W202–208 (2009). 10.1093/nar/gkp335

36 Hartmann, M. et al. Molecular Plasticity Results in Oncofetal Reprogramming and Therapeutic Vulnerabilities in Juvenile Myelomonocytic Leukemia. Blood Cancer Discov 7, 306–330 (2026). 10.1158/2643-3230.BCD-25-0246

37 Louka, E. et al. Heterogeneous disease-propagating stem cells in juvenile myelomonocytic leukemia. J Exp Med 218 (2021). 10.1084/jem.20180853

38 Aparicio-Puerta, E. et al. sRNAbench and sRNAtoolbox 2022 update: accurate miRNA and sncRNA profiling for model and non-model organisms. Nucleic Acids Res 50, W710–W717 (2022). 10.1093/nar/gkac363

39 Love, M. I., Huber, W. & Anders, S. Moderated estimation of fold change and dispersion for RNA-seq data with DESeq2. Genome biology 15, 550 (2014). 10.1186/s13059-014-0550-8

40 Gu, Z. Complex heatmap visualization. Imeta 1, e43 (2022). 10.1002/imt2.43

41 Wilbert, M. L. et al. LIN28 binds messenger RNAs at GGAGA motifs and regulates splicing factor abundance. Mol Cell 48, 195–206 (2012). 10.1016/j.molcel.2012.08.004

42 Hafner, M. et al. Identification of mRNAs bound and regulated by human LIN28 proteins and molecular requirements for RNA recognition. RNA 19, 613–626 (2013). 10.1261/rna.036491.112

43 Tao, T. et al. LIN28B regulates transcription and potentiates MYCN-induced neuroblastoma through binding to ZNF143 at target gene promotors. Proceedings of the National Academy of Sciences of the United States of America 117, 16516–16526 (2020). 10.1073/pnas.1922692117

44 Hovanes, K. et al. Beta-catenin-sensitive isoforms of lymphoid enhancer factor-1 are selectively expressed in colon cancer. Nat Genet 28, 53–57 (2001). 10.1038/88264

45 Schonung, M. et al. International Consensus Definition of DNA Methylation Subgroups in Juvenile Myelomonocytic Leukemia. Clin Cancer Res 27, 158–168 (2021). 10.1158/1078-0432.CCR-20-3184

46 Shi, J. et al. LIN28B inhibition sensitizes cells to p53-restoring PPI therapy through unleashed translational suppression. Oncogenesis 11, 37 (2022). 10.1038/s41389-022-00412-8

47 Wander, P. et al. High-throughput drug screening reveals Pyrvinium pamoate as effective candidate against pediatric MLL-rearranged acute myeloid leukemia. Transl Oncol 14, 101048 (2021). 10.1016/j.tranon.2021.101048

48 Yao, K. et al. Wnt Regulates Proliferation and Neurogenic Potential of Muller Glial Cells via a Lin28/let-7 miRNA-Dependent Pathway in Adult Mammalian Retinas. Cell Rep 17, 165–178 (2016). 10.1016/j.celrep.2016.08.078

49 Luo, Y. et al. The osteogenic differentiation of human adipose-derived stem cells is regulated through the let-7i-3p/LEF1/beta-catenin axis under cyclic strain. Stem Cell Res Ther 10, 339 (2019). 10.1186/s13287-019-1470-z

50 Rodriguez-Ubreva, J. et al. C/EBPa-mediated activation of microRNAs 34a and 223 inhibits Lef1 expression to achieve efficient reprogramming into macrophages. Mol Cell Biol 34, 1145–1157 (2014). 10.1128/MCB.01487-13

51 Liang, J. et al. LEF1 targeting EMT in prostate cancer invasion is mediated by miR-181a. Am J Cancer Res 5, 1124–1132 (2015).

52 Igarashi, K. J. et al. Adult hematopoietic stem cells activate a normally fetal-restricted program to functionally expand ex vivo. Blood (2026). 10.1182/blood.2025030268

53 Eldeeb, M. et al. A fetal tumor suppressor axis abrogates MLL-fusion-driven acute myeloid leukemia. Cell Rep 42, 112099 (2023). 10.1016/j.celrep.2023.112099

54 Okeyo-Owuor, T. et al. The efficiency of murine MLL-ENL-driven leukemia initiation changes with age and peaks during neonatal development. Blood Adv 3, 2388–2399 (2019). 10.1182/bloodadvances.2019000554

55 Li, Y. et al. LIN28B promotes differentiation of fully transformed AML cells but is dispensable for fetal leukemia suppression. Leukemia 38, 648–651 (2024). 10.1038/s41375-024-02167-0

56 Zhou, J. et al. Inhibition of LIN28B impairs leukemia cell growth and metabolism in acute myeloid leukemia. J Hematol Oncol 10, 138 (2017). 10.1186/s13045-017-0507-y

57 Zhou, J. et al. LIN28B Activation by PRL-3 Promotes Leukemogenesis and a Stem Cell-like Transcriptional Program in AML. Molecular cancer research : MCR 15, 294–303 (2017). 10.1158/1541-7786.MCR-16-0275-T

58 Zhou, M. et al. RNA-Binding Protein Lin28B Promotes Chronic Myeloid Leukemia Blast Crisis by Transcriptionally Upregulating miR-181d. Molecular cancer research : MCR 22, 932–942 (2024). 10.1158/1541-7786.MCR-23-0928

59 Chen, Y. et al. LIN28/let-7/PD-L1 Pathway as a Target for Cancer Immunotherapy. Cancer Immunol Res 7, 487–497 (2019). 10.1158/2326-6066.CIR-18-0331

60 Matias-Barrios, V. M. et al. Developing novel Lin28 inhibitors by computer aided drug design. Cell Death Discov 11, 5 (2025). 10.1038/s41420-024-02281-z

61 Schultz, C. W. & Nevler, A. Pyrvinium Pamoate: Past, Present, and Future as an Anti-Cancer Drug. Biomedicines 10 (2022). 10.3390/biomedicines10123249

