## Supplementary information for "LIN28B is a post-transcriptional regulator of Wnt signalling in myeloid leukemia cells and dual targeting promotes cytotoxicity"

### Supplementary methods

*Cell culture and drug treatments*

The myeloid cell lines are used in this study are as follows: K562, HL60, HEL, U937, PLB-985, NOMO-1, OCI-AML3, EOL-1, ML-1, THP-1, KU812 (The European Collection of Authenticated Cell Cultures) and OCI-AML2, KG-1, KG1a, SET-2, NB4 and MONO-MAC6 (Leibniz Institute DSMZ-German Collection of Microorganisms and Cell Cultures GmbH). Short-tandem repeat (STR) analysis was done for cell line authentication for these myeloid cell cultures. Myeloid cell lines were cultured in RPMI-1640 medium, supplemented with L-glutamine and streptomycin at 37°C with 5% $CO_2$. HEK293T cell line was utilized for lentivirus preparation and cultured in DMEM supplemented with L-glutamine and streptomycin at 37°C with 5% $CO_2$. β-Catenin stabilisation was achieved through the GSK-3β inhibitor CHIR99021 (Merk-Millipore, Gillingham, Dorset) as previously described.[1] LIN28-let-7 interaction was globally blocked using compound 1632 (C1632)[2] (SML3673-10MG, Merck-Millipore). Six different inhibitors (BC2059 (MedChemExpress [MCE], HY-109103), MSAB (MCE, HY-120697), KYA1797K (MCE, HY-101090), LGK974 (MCE, HY-17545), Pyrvinium Pamoate (MCE, HY-A0293), XAV-939 (MCE, HY-15147)) were tested to achieve pharmacological inhibition of β-catenin expression in myeloid cell lines.

*Primary samples*

Human cord blood (CB) was obtained following informed consent from healthy mothers at full-term undergoing elective caesarean sections at from University Hospitals Sussex and East Sussex Healthcare NHS Foundation Trusts with approval the East of England-Essex Research Ethics Committee, HRA and Health and Care Research Wales (18/EE/0403). Mononuclear cells (MNCs) with viability >80% following isolation via density gradient separation using Ficoll-Hypaque (Merck-Millipore, Gillingham, Dorset) were included in the study and cryopreserved in liquid nitrogen until experimental use.

Donated fetal tissue was provided for purposes of this research by the Human Developmental Biology Resource (www.hdbr.org), regulated by the UK Human Tissue Authority (www.hta.gov.uk) and covered under ethics granted by NHS Health Research Authorities: North East—Newcastle & North Tyneside Research Ethics

Committee (REC: 18/NE/0290) and London—Fulham Research Ethics Committee (18/LO/0822). Informed consent was obtained from all participants, who donated human fetal tissue for research without receiving any monetary compensation. The $CD34^+$ HSPC fraction of human cord blood and fetal livers was enriched to >80% purity as previously described[3] using MiniMACS CD34 microbeads (Miltenyi Biotec, Woking, Surrey) according to the manufacturer's instructions and confirmed by flow cytometric assessment of CD34 positivity.

*Lentivirus preparation and transduction*

MISSION lentiviral shRNA plasmids and lentiviral overexpression plasmids were obtained from Merck-Millipore and VectorBuilder, respectively (see Supplemental Table S4 for details). Plasmid DNAs were supplied as pre-transformed glycerol stocks, which were streaked onto agar plates to isolate single colonies. Colonies were incubated overnight at 37°C, and plasmid DNA was purified using the GeneJET Plasmid Miniprep Kit (Thermo Fisher Scientific, Altrincham, Cheshire). HEK293T cells were used to produce lentivirus. Tissue culture–treated flasks were coated with 0.01% poly-L-lysine solution (Merck Millipore), after which 4–5 x $10^6$ HEK293T cells were seeded to achieve ~70% confluency the following day. Lentiviral particles were generated using Lipofectamine 3000 Transfection Reagent (Thermo Fisher Scientific), Opti-MEM I (1X) (Thermo Fisher Scientific), and the appropriate envelope, packaging, and transfer plasmids. Viral supernatants were harvested 24- and 48-hours post-transfection. For transduction, myeloid cell lines were plated in non-treated 24-well tissue culture plates (Thermo Fisher Scientific). Plates were first coated with 40 µg/ml Retronectin (Takara Bio, London, UK) overnight at 4°C, then blocked with 1% BSA in PBS for 30 minutes. Virus-containing supernatants were centrifuged at 2000 × g for 90 minutes at room temperature to promote viral sedimentation. Subsequently, 1 ml of resuspended myeloid cells (2–5x$10^5$) was added per well. Following 2 days of viral integration and cell expansion, cells were subjected to drug selection with 1 µg/ml puromycin using the corresponding parental cell line as control. Protein knockdown or overexpression was confirmed by immunoblotting.

*Immunoblotting*

Whole-cell lysates were prepared using ice-cold 10x cell lysis buffer (20 mM Tris-HCl, pH 7.5; 150 mM NaCl; 1 mM EGTA; 1% Triton; 2.5 mM sodium pyrophosphate; 1 mM β-glycerophosphate; 1 mM $Na_3VO_4$; 1 µg/mL leupeptin - Cell Signaling Technology,

Leiden, Netherlands) diluted with distilled water and supplemented with one tablet of Complete Mini Protease Inhibitor (Roche, Welwyn Garden City, Hertfordshire). Protein concentrations were determined using the Detergent-Compatible Colorimetric Assay Kit (BioRad) to ensure equal protein loading per sample. Lysates were diluted with 1x Laemmli buffer (Thermo Fisher Scientific) and distilled water, then heated at 95 °C for 5 minutes before immunoblotting. SDS-PAGE (sodium dodecyl sulfate–polyacrylamide gel electrophoresis) was performed to separate proteins, which were subsequently transferred onto Immobilon-P Hydrophobic PVDF Transfer Membranes (Merck-Millipore). Gels ranging from 7.5% to 15% acrylamide were used depending on the size of the protein of interest. Membranes were probed overnight at 4 °C with antibodies against the target proteins. Primary antibodies used in this study are listed in Supplemental Table S5. Target proteins were visualized using the LumiGLO® Peroxidase Chemiluminescent Substrate Kit (Sera Care, Wembley, UK).

*Cytoplasmic/nuclear fractionation*

A total of $5x10^6$ cells were harvested for cytoplasmic and nuclear fractionation. After washing the cells with ice-cold PBS, each pellet was resuspended in cytoplasmic lysis buffer (10mM Tris-HCl; pH 8, 10mM NaCl, 1.5mM $MgCl_2$, and 0.5% Igepal-CA630/NP40; supplemented with one tablet of Complete Mini Protease Inhibitor (Roche)). The samples were centrifuged at 800 x g for 5 minutes at 4°C, and the supernatant was collected as the cytoplasmic fraction. The pellet was then washed with ice-cold PBS, resuspended in nuclear lysis buffer (Cell Signalling Technology), and subjected to sonication. Following sonication, the nuclear lysates were centrifuged at 17,000 × g for 10 minutes at 4°C, and the supernatant was collected as the nuclear fraction. Both cytoplasmic and nuclear fractions were quantified and processed for immunoblotting as described for whole-cell lysates above.

*Co-immunoprecipitation (Co-IP)*

Protein G Dynabeads (Thermo Fisher Scientific) were coupled with 8 µg of LIN28B antibody (Cell Signalling Technology, 4196) or rabbit IgG (Cell Signalling Technology, 2729). The beads were crosslinked to the antibodies using 20 mM dimethyl pimelimidate. Next, 1 mg of whole-cell lysate was incubated overnight at 4 °C with the crosslinked bead–antibody complexes in Co-IP lysis buffer (20 mM Tris-HCl, pH 7.5; 150 mM NaCl; 1 mM EGTA; 1% Triton; 2.5 mM sodium pyrophosphate; 1 mM β-glycerophosphate; 1 mM $Na_3VO_4$; 1 µg/mL leupeptin; 10% glycerol; and 0.5 mM DTT). After incubation, the samples were washed with Co-IP wash buffer (phosphate-buffered saline (PBS) containing 0.02% Tween-20). Each sample was then diluted with 2× Laemmli buffer (Thermo Fisher Scientific) and Co-IP lysis buffer, and incubated at 95°C for 5 minutes before immunoblotting. 20 µg/ml RNase A (Thermo Fisher Scientific) was added to the Co-IP lysates to remove RNA molecules.

*RNA extraction and clean-up*

Total RNA extraction was done using Zymo Research Quick-RNA™ Miniprep Kit (Cambridge Bioscience, Cambridge, Cambridgeshire) and the extracted total RNA was exposed to DNase treatment to remove genomic DNA from RNA samples through TURBO DNA-free™ Kit (Thermo Fisher Scientific). Manufacturer's guidelines were followed.

*RT-qPCR*

The abundance of target transcripts was detected using One-Step RT-qPCR Mix (APTO-GEN, London, UK). The RT-qPCR reactions was prepared as follows: RT+ reaction (5 µl 2x SyGr RT-qPCR mix, 0.5 µl 20x reverse transcriptase, 0.25 µl forward/reverse primer, 1 µl 10ng of RNA, 3 µl nuclease-free water) and RT- reaction (5 µl 2x SyGr RT-qPCR mix, 0.25 µl forward/reverse primer, 1 µl 10ng of RNA, 3.5 µl nuclease-free water). RT-qPCR was performed under the recommended thermocycler conditions: 1 cycle (50°C, 10 minutes), 1 cycle (95°C, 1 minutes) and 40 cycles (95°C, 5 seconds & 60°C 20 seconds). Forward and reverse primer sequences used in this study are listed in Supplemental Table S5 below and were optimised to show efficiency between 90-110%.

*Primary JMML scRNA sequencing analysis*

- Re-analysis of scRNA-seq data of primary JMML samples from Hartmann et al[4]

Because *LIN28B* and *LEF1* transcripts were sparsely detected in the single-cell RNA-seq dataset, likely reflecting both biological heterogeneity and the limited sensitivity of droplet-based scRNA-seq, complementary analytical approaches were employed to evaluate potential relationships between *LIN28B* and *LEF1*. These included (i) co-detection enrichment analyses based on binary transcript detection, (ii) expression correlation analyses restricted to double-positive cells, and (iii) module score analyses of associated transcriptional programs.

*LIN28B/LEF1 co-detection enrichment analysis*: Individual cells were classified as *LIN28B*-positive or *LEF1*-positive according to transcript detection (normalized expression >0). For each subgroup, the expected number of *LIN28B+*/*LEF1+* double-positive cells under the assumption of independent transcript detection was calculated as:

E = (N(*LIN28B*+) x N(*LEF1*+)) / N(total),

where N(*LIN28B*+) and N(*LEF1*+) denote the numbers of *LIN28B*-positive and *LEF1*-positive cells and N(total) denotes the total number of analysed cells in the respective subgroup. Co-detection enrichment was quantified as the ratio of observed to expected double-positive cells (observed/expected ratio). Statistical significance of deviation from independence was assessed using two-sided Fisher's exact tests applied to 2×2 contingency tables of *LIN28B* and *LEF1* detection status.

*Co-expression analysis of LIN28B/LEF1 double-positive cells*: Expression concordance between *LIN28B* and *LEF1* was assessed in HSPCs, the proposed cellular compartment of origin in JMML. Correlation analyses were restricted to cells with detectable expression of both transcripts (expression >0 for both genes) to minimise the influence of sparse transcript detection and zero inflation. Associations between *LIN28B* and *LEF1* expression levels were quantified using two-sided Spearman rank correlation and visualised by least-squares regression lines. Fetal HSPCs were included as a developmental reference population because *LIN28B* expression has previously been shown in this compartment.

*LIN28B- and LEF1-associated transcriptional programs*: To assess coordinated transcriptional states beyond individual transcript detection, *LIN28B*- and *LEF1*-associated gene programs were derived from normal reference HSPCs ($CD34^+$ HSC,

CD34$^+$ HSC-cycle and CD34$^+$ MultiLin cells from fetal, neonatal, juvenile and adult samples). For each anchor gene, Spearman correlations were calculated between normalized anchor-gene expression and all genes detected in at least 1% of reference HSPCs. The 50 most positively correlated genes were retained after exclusion of the anchor gene itself. Per-cell module scores were calculated across the complete dataset using Seurat AddModuleScore. Associations between *LIN28B* and *LEF1* module scores were evaluated across normal and JMML HSPCs using Spearman correlation analysis.

- Re-analysis of scRNA-seq data of primary JMML samples from Louka et al.[5]

To examine the correlation between LIN28B and Wnt/β-catenin signalling, a publicly available dataset of JMML patient samples (GSE111709) was used. After raw data were obtained, preprocessing was performed using fastp for trimming, HISAT2 for alignment to the human genome, and featureCounts for counting the mapped reads. Raw counts were normalised using edgeR in R software, and the normalised counts were then used for visualisation and the heatmap.

*miRNA sequencing*

2x10$^6$ cells were harvested for each of the three biological replicates per condition (β-catenin shRNA, LIN28B shRNA and non-targeting shRNA). Total RNA extraction was performed using the Quick-RNA™ Miniprep Kit (Cambridge Bioscience) as described in manufacturer's guidelines. Quality control of total RNA samples was done with the Agilent TapeStation 4150. Lastly, the RNA samples were sent for miRNA sequencing (miRNA-seq) to BGI Genomics (Shenzhen, China) for library preparation and sequencing.

BGI Genomics provided clean FASTQ files, and we pre-processed them in sRNAbench on the sRNAtoolbox platform,[6] with the following parameters: miRbase release 22.1 (as miRNA annotation reference database), Homo sapiens (hsa) (as short name from miRbase), human (GRCh38_p13) (as species), "Provided reads are already trimmed" (as sequencing library protocol), the rest of parameters as defaults. Raw read counts obtained after preprocessing was used to determine upregulated or downregulated miRNAs with β-catenin or LIN28B knockdown versus control using

DESeq2 (version 1.44.0) package in R software.[7] Also, tidyverse (version 2.0.0) and ComplexHeatmap (version 2.20.0) packages were used for data manipulation and visualisation.[8]

*RNA stability assay*

K562 cells were seeded at a density of 1x$10^6$ cells/ml, and 5 µg/ml Actinomycin D (Merck Millipore) was added to the cells. Samples were collected at 0-, 4- and 8-hour time points, and RNA extraction, DNase treatment and RT-qPCR was performed as described above. The RNA stability of the target RNAs were calculated based on the RNA abundance in 0-time point for each cell line separately.

*Flow cytometry*

Wnt/β-catenin signalling activity was measured in 2x$10^5$ myeloid cells using the BAR/fuBAR reporter system.[9] Cells were treated overnight with either DMSO or CHIR99021. TCF/LEF activity was assessed by flow cytometric analysis of Venus yellow fluorescent protein (YFP) intensity. Cell viability of myeloid cells with β-catenin knockdown after C1632 treatment was assessed using 7-AAD viability staining solution (Thermo Fisher Scientific), whereas cell viability was evaluated using the Zombie Violet Fixable Viability Kit (BioLegend, London, UK) when the β-catenin inhibitor pyrvinium pamoate was used. This was because pyrvinium pamoate exhibits red fluorescence, which interferes with the 7-AAD staining channel during flow cytometry. Data acquisition was performed on a CytoFLEX flow cytometer (Beckman Coulter, California, USA), with a minimum of 2x$10^4$ debris-excluded events collected per sample. Flow cytometric data were analysed using FlowJo software (Tree Star Inc., Ashland, OR, USA).

**Supplemental Table S1:** Lentivirus plasmids used for transgene expression in this study.

| Target gene | Expression type | Vector type | Supplier |
|---|---|---|---|
| Non-targeting | shRNA control | pLKO.1-puro Non-Mammalian shRNA Control Plasmid SHC002 | Merck MISSON® |
| *LIN28B* | shRNA (#1) | pLKO_TRCN0000122191 | Merck MISSON® |
| *LIN28B* | shRNA (#2) | pLKO_TRCN0000122599 | Merck MISSON® |
| *CTNNB1* | shRNA (#1) | pLKO_TRCN0000314920 | Merck MISSON® |
| *CTNNB1* | shRNA (#2) | pLKO_TRCN0000314921 | Merck MISSON® |
| Empty vector control | Ectopic | pLV[Exp]-Puro-EF1A>ORF_91bp (VB160723-1006snj) | VectorBuilder |
| *LIN28B* | Ectopic | pLV[Exp]-Puro-EF1A>hLIN28B[NM_001410939.1] (VB230202-1438syb) | VectorBuilder |

**Supplemental Table S2:** Primary antibodies used for immunoblotting in this study.

| Antibody | Source |
|---|---|
| β-Catenin | BD Biosciences, 610153 |
| LIN28B | Cell Signalling Tech, 4196 |
| LEF-1 | Cell Signalling Tech, 2230 |
| TCF4 (TCF7L2) | Cell Signalling Tech, 2569 |
| Lamin A/C | Merck Millipore, SAB4200236 |
| α-Tubulin | Merck Millipore, T9026 |
| GAPDH | Proteintech, 6004-1-Ig |
| LIN28A | Cell Signalling Tech, 3978 |
| β-Actin | Merck Millipore, A1978 |

**Supplemental Table S3:** Forward and reverse primer sequences used for RT-qPCR in this study.

| Target | Forward Primer (5\` → 3\`) | Reverse Primer (5\` → 3\`) |
|---|---|---|
| *LIN28B* | CATCTCCATGATAAACCGAGAGG | GTTACCCGTATTGACTCAAGGC |
| *LEF1* | AGAACACCCCGATGACGGA | GGCATCATTATGTACCCGGAAT |
| *TCF7L2* | GCCTCTTATCACGTACAGCAAT | GCCAGGCGATAGTGGGTAAT |
| *GAPDH* | GCATCTTCTTTTGCGTCGCC | CCCAATACGACCAAATCCGTTG |
| *MYC* | TCCCTCCACTCGGAAGGAC | CTGGTGCATTTTCGGTTGTTG |
| *RNA18S* | CTCAACACGGGAAACCTCAC | CGCTCCACCAACTAAGAACG |
| *LEF1 (Intronic Primers)* | AGCACAAACCTCTCAGGAGC | ACAAATTGCACCCCTTATCTGC |

### Supplementary Figures

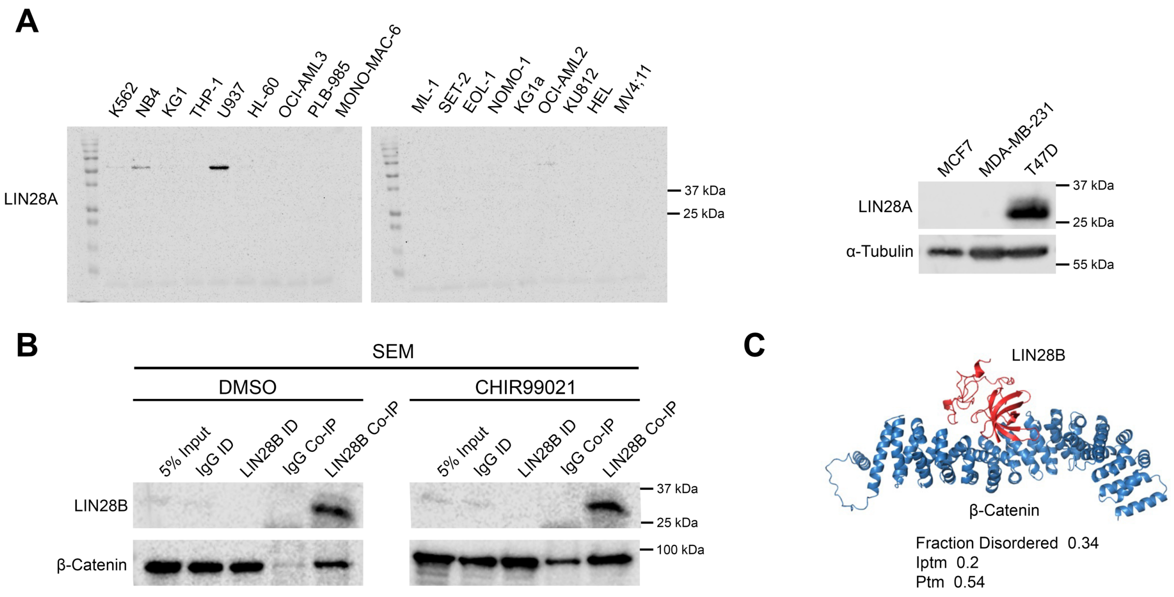


**Supplementary Figure S1. (A)** Immunoblots showing absence of LIN28A protein expression across 18 myeloid leukaemia cell lines (same as main Figure 1C) and three breast cancer cell lines. α-Tubulin served as a loading control. **(B)** Immunoblots showing β-catenin levels in LIN28B co-IPs from SEM whole-cell lysates following overnight treatment with ± 5 μM CHIR99021. **(C)** AlphaFold 3–predicted model of the β-catenin (blue)–LIN28B (red) complex.

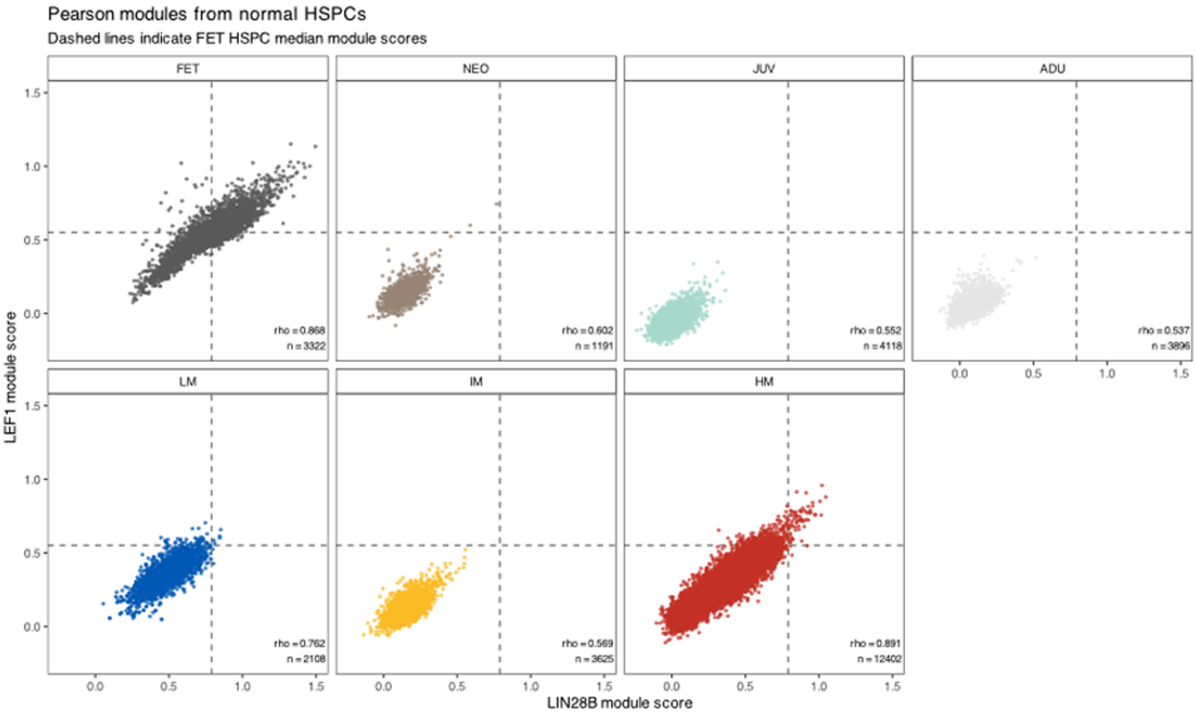


**Supplementary Figure S2.** Subgroup-specific correlations of normal HSPC-derived *LIN28B* and *LEF1* module scores. Individual HSPCs are shown separately for each developmental and JMML subgroup. Dashed lines indicate the median *LIN28B* and *LEF1* module scores of fetal HSPCs. Spearman correlation coefficients (ρ) and numbers of analysed cells (n) are indicated.

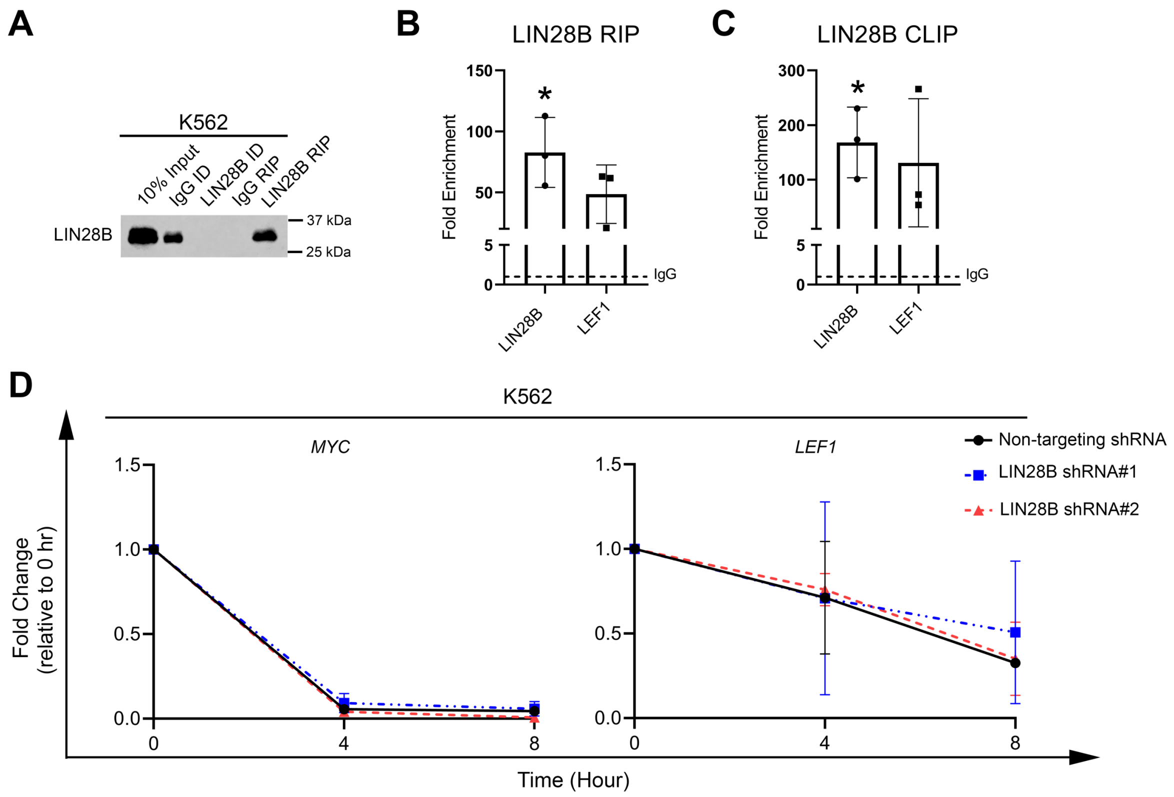


**Supplementary Figure S3. (A)** Immunoblot showing the enrichment of LIN28B protein in LIN28B RIP samples from K562 cells. Bar graphs show the fold enrichment of *LIN28B* and *LEF1* relative to IgG controls (dashed) following **(B)** RIP- or **(C)** CLIP-RT-qPCR in K562 cells. **(D)** Line graphs summarizing the fold change in *MYC* and *LEF1* levels in K562 cells with ± LIN28B shRNA following Actinomycin D treatment. *RNA18S* was used to normalize total RNA levels, and normalization for each condition was performed by comparing the abundance of target mRNA at each time point to the respective 0-hour level for each cell line. Data are presented as mean ± SD from three independent biological replicates. Statistical significance was assessed using Student's t-test (*$p < 0.05$).

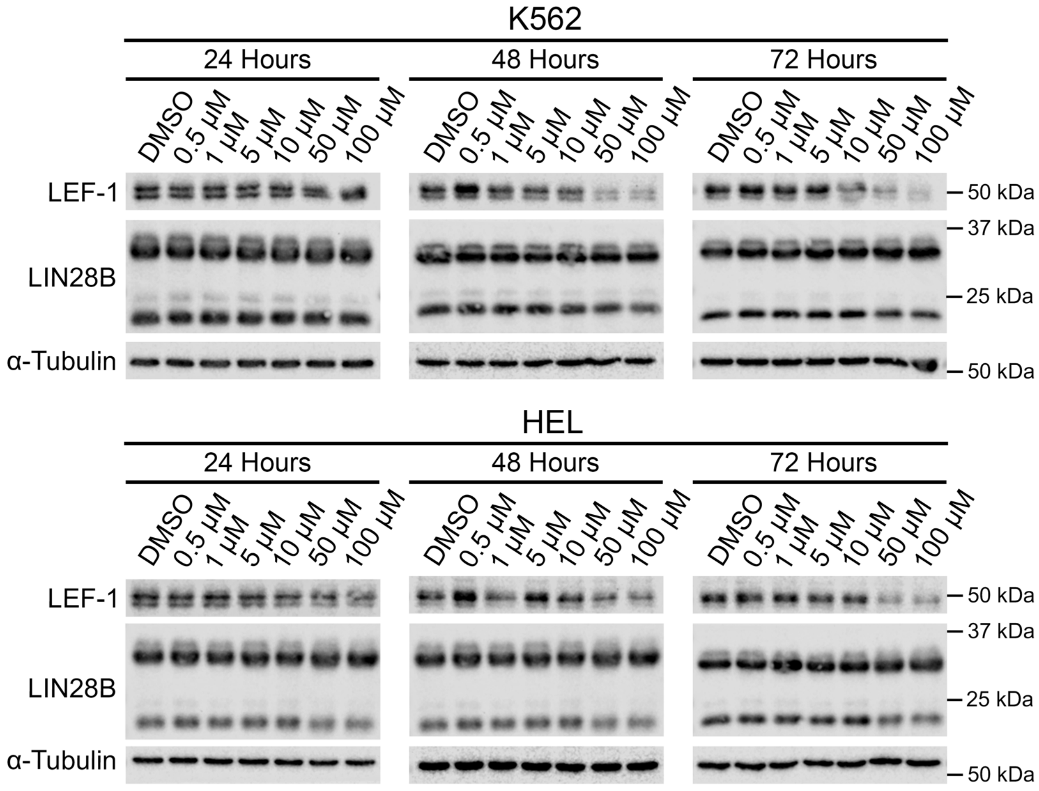


**Supplementary Figure S4.** Immunoblots showing expression of LIN28B and LEF-1 protein in K562 and HEL cells exposed to 0.5-100µM of the LIN28B:*let7* inhibitor ln268. α-Tubulin served as a loading control.

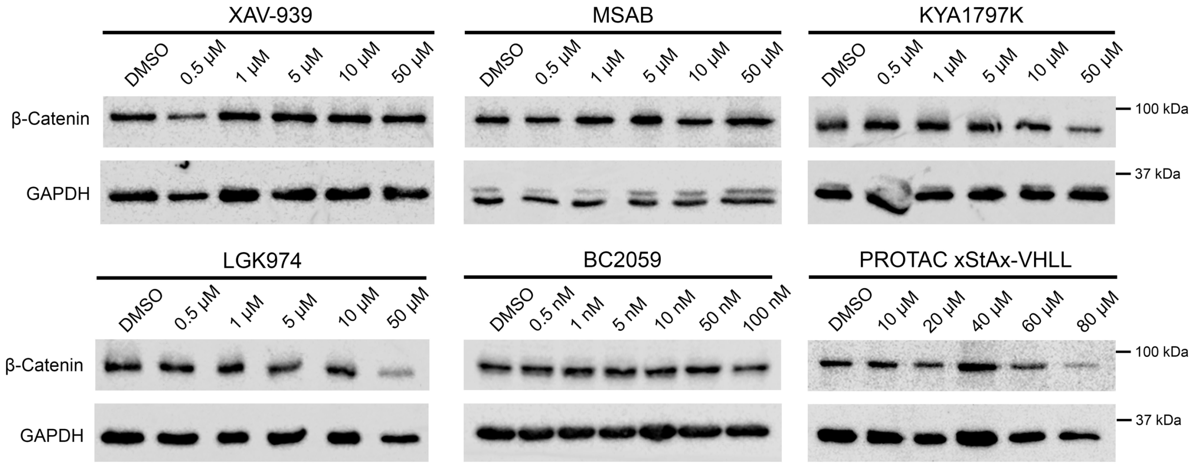


**Supplementary Figure S5.** Immunoblots showing β-catenin protein expression levels in HEL cells following overnight treatment with different β-catenin inhibitors. BC2059 was used in nanomolar range due to excessive cytotoxicity in micromolar testing. GAPDH was used as a loading control.

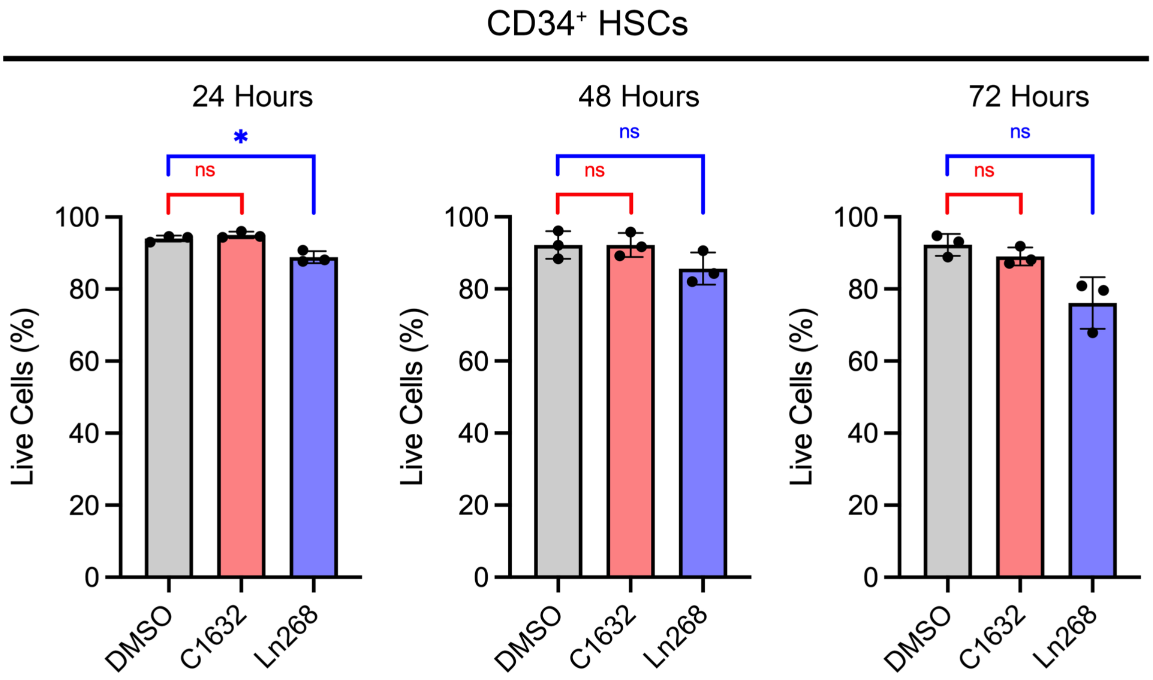


**Supplementary Figure S6. (A)** Bar graphs showing the percentage viability of human CB-derived CD34+ HSPCs treated with DMSO vehicle control, 100μM C1632 or 100μM Ln268 for 24, 48 and 72hrs, as determined by negative 7AAD staining via flow cytometry. Data are presented as mean ± SD from three independent biological replicates. Statistical significance was assessed using Student's t-test (*$p < 0.05$; ns: non-significant).

**Supplementary Table S4:** Upregulated and downregulated miRNAs following β-catenin knockdown in K562 cells.

| **β-Catenin Knockdown** | | | |
|---|---|---|---|
| **Upregulated miRNAs** | **Downregulated miRNAs** | | |
| hsa-miR-1301-3p | hsa-miR-520a-5p | hsa-miR-516b-5p | hsa-miR-5189-3p |
| hsa-miR-181c-5p | hsa-miR-671-3p | hsa-miR-3689a-5p | hsa-miR-431-5p |
| hsa-miR-181c-3p | hsa-miR-518b | hsa-miR-944 | hsa-miR-487b-3p |
| hsa-miR-488-3p | hsa-miR-512-3p | hsa-miR-498-5p | hsa-miR-520h |
| hsa-miR-181d-5p | hsa-miR-516a-5p | hsa-miR-520f-3p | hsa-miR-4772-3p |
| hsa-miR-935 | hsa-miR-411-3p | hsa-miR-517-5p | hsa-miR-520g-3p |
| hsa-miR-1270 | hsa-miR-520e-3p | hsa-miR-3689e | hsa-miR-382-5p |
| hsa-miR-193a-3p | hsa-miR-3940-3p | hsa-miR-363-3p | hsa-miR-519d-3p |
| hsa-miR-210-3p | hsa-miR-494-5p | hsa-miR-376b-3p | hsa-miR-889-3p |
| hsa-miR-34a-5p | hsa-miR-409-3p | hsa-miR-520c-3p | hsa-miR-4435 |
| hsa-miR-100-5p | hsa-miR-212-5p | hsa-miR-523-3p | hsa-miR-410-3p |
| hsa-miR-373-3p | hsa-miR-376a-3p | hsa-miR-518a-3p | hsa-miR-3681-5p |
| hsa-miR-193b-3p | hsa-miR-369-5p | hsa-miR-517b-3p | hsa-miR-337-3p |
| hsa-miR-449b-5p | hsa-miR-517a-3p | hsa-miR-3689b-5p | |

**Supplementary Table S5:** Commonly regulated miRNAs ($|\log_2FC| > 1$ and $p$-adj $< 0.05$) following β-catenin and LIN28B knockdown in K562 cells. (K/d: Knockdown)

| Upregulated miRNAs | Downregulated miRNAs |
|---|---|
| hsa-miR-1301-3p | hsa-miR-4772-3p |
| hsa-miR-488-3p | hsa-miR-3681-5p |
| hsa-miR-935 | |
| hsa-miR-193a-3p | |
| hsa-miR-100-5p | |
| hsa-miR-373-3p | |
| hsa-miR-449b-5p | |

| Downregulated in β-catenin K/d but upregulated in LIN28B K/d miRNAs | | |
|---|---|---|
| hsa-miR-520a-5p | hsa-miR-517-5p | hsa-miR-3689a-5p |
| hsa-miR-518b | hsa-miR-3689e | hsa-miR-944 |
| hsa-miR-512-3p | hsa-miR-520c-3p | hsa-miR-498-5p |
| hsa-miR-516a-5p | hsa-miR-523-3p | hsa-miR-520f-3p |
| hsa-miR-520e-3p | hsa-miR-518a-3p | hsa-miR-520h |
| hsa-miR-517a-3p | hsa-miR-517b-3p | hsa-miR-520g-3p |
| hsa-miR-516b-5p | hsa-miR-3689b-5p | hsa-miR-519d-3p |

### References


1 Morgan, R. G. *et al.* LEF-1 drives aberrant beta-catenin nuclear localization in myeloid leukemia cells. *Haematologica* **104**, 1365-1377 (2019). https://doi.org:10.3324/haematol.2018.202846

2 Roos, M. *et al.* A Small-Molecule Inhibitor of Lin28. *ACS Chem Biol* **11**, 2773-2781 (2016). https://doi.org:10.1021/acschembio.6b00232

3 Morgan, R. G. *et al.* gamma-Catenin is expressed throughout normal human hematopoietic development and is required for normal PU.1-dependent monocyte differentiation. *Leukemia* **27**, 2096-2100 (2013). https://doi.org:10.1038/leu.2013.96

4 Hartmann, M. *et al.* Molecular Plasticity Results in Oncofetal Reprogramming and Therapeutic Vulnerabilities in Juvenile Myelomonocytic Leukemia. *Blood Cancer Discov* **7**, 306-330 (2026). https://doi.org:10.1158/2643-3230.BCD-25-0246

5 Louka, E. *et al.* Heterogeneous disease-propagating stem cells in juvenile myelomonocytic leukemia. *J Exp Med* **218** (2021). https://doi.org:10.1084/jem.20180853

6 Aparicio-Puerta, E. *et al.* sRNAbench and sRNAtoolbox 2022 update: accurate miRNA and sncRNA profiling for model and non-model organisms. *Nucleic Acids Res* **50**, W710-W717 (2022). https://doi.org:10.1093/nar/gkac363
7 Love, M. I., Huber, W. & Anders, S. Moderated estimation of fold change and dispersion for RNA-seq data with DESeq2. *Genome biology* **15**, 550 (2014). https://doi.org:10.1186/s13059-014-0550-8
8 Gu, Z. Complex heatmap visualization. *Imeta* **1**, e43 (2022). https://doi.org:10.1002/imt2.43
9 Biechele, T. L. & Moon, R. T. Assaying beta-catenin/TCF transcription with beta-catenin/TCF transcription-based reporter constructs. *Methods Mol.Biol.* **468**, 99-110 (2008).
